# Adherent reformed islets: a long-term primary cell-based platform for exploring mouse and human islet biology

**DOI:** 10.1101/2023.11.22.568245

**Authors:** N. Haq, K.W. Toczyska, M.E. Wilson, M. Jacobs, Min Zhao, Y. Lei, J. Pearson, S.J. Persaud, T.J. Pullen, G.A. Bewick

## Abstract

Pancreatic islets are 3-dimensional micro-organs that maintain β-cell functionality via cell-cell and cell-matrix communication. Isolated primary islets are the gold standard for in vitro models. However, native islets present experimental challenges for long-term mechanistic studies owing to their short culture life (approximately 1 week). We developed a novel long-term protocol to study the function of primary islets. The protocol employed reformed islets following dispersion and a fine-tuned culture environment. Reformed islets are highly similar to their primary counterparts across various physiological characteristics. Long-term culture of reformed islets enables high-resolution imaging, repeated functional assessment, and the study of cell-cell communication. Unlike other platforms such as stem cell-derived organoids, reformed islets retain their resident immune populations, making them ideal for studying both resident and infiltrating immune cells and their interactions with hormone-producing islet cells.

Qualitative and quantitative analyses revealed that the composition and cytoarchitecture of the reformed islets mimicked those found in primary islets, including the presence of macrophages and CD4^+^ and CD8^+^ T cells, which are the key resident immune cell types. Reformed islets secrete insulin and are glucose-responsive, and their β-cells can be stimulated to proliferate using GLP-1 receptor agonism. Furthermore, a comparison of the transcriptomic landscape of isolated human islets and reformed islets generated from the same donor demonstrated a high degree of similarity.

Our reformed islets provide an ideal platform to study diabetes pathology. We recapitulated both the T1DM and T2DM disease milieu and validated our model for studying islet immune trafficking and invasion using activated macrophages and T cells.

Our data illustrates that reformed islets are an anatomical and functional alternative to native human and mouse islets. Moreover, reformed islets have an advantage over mouse and human β-cell lines, including MIN6 and EndoC-βH1cells, that lack the signalling input of non-β-endocrine cells and immune cell crosstalk. In this study, we showed that reformed islets are a durable paradigm (cell-based model) for islet-based exploration and a means of target discovery/validation for diabetes research.

**Graphical Abstract:** 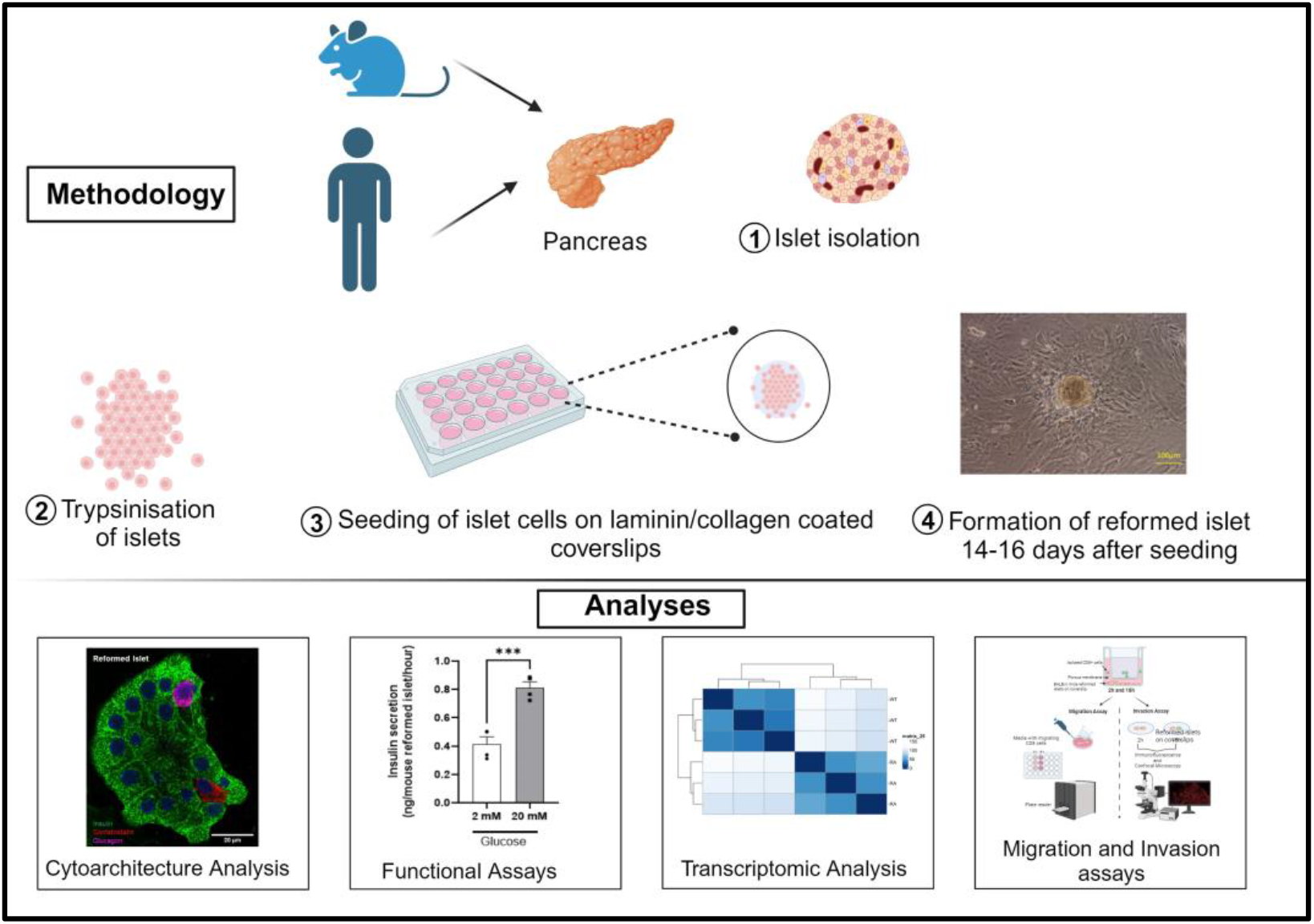

## Introduction

The prevalence of both type 1 and type 2 diabetes mellitus (T1DM, T2DM), which are among the leading causes of morbidity and mortality, is rapidly increasing, making it one of the largest global health concerns, affecting 537 million adults worldwide [1-3]. Although effective treatments exist for T2DM, intensive insulin therapy and islet transplantation for a limited number of people are the only approved option for T1DM despite improvements in care. For both types, treatment aims to manage symptoms and reduce complications; however, there is no cure.

Diabetes pathogenesis involves a complex interplay of multiple mechanisms, including genetics, islet/β-cell dysfunction, and immune dysregulation. As such, efforts to understand disease pathology have focused on comprehensively exploring and understanding all aspects of islet biology. Islets are micro-organs that regulate glucose homeostasis through the secretion of the key hormones insulin, glucagon, and somatostatin. They comprise interspersed β, α, δ, ε, and PP cells intermingled with resident immune cells, stellate cells, smooth muscles, and fibroblasts [4]. β-cell functionality is maintained via cell-cell and cell-matrix communication. Therefore, understanding islet physiology and pathological dysfunction requires exquisite knowledge of how all islet cells communicate to maintain normal function, and how this is dysregulated to drive disease progression.

Primary islets, often isolated from rodents, are the gold standard for diabetes research. They are readily available and cost-effective, and their function is comparable to those observed *in vivo* [5]. However, isolating pancreatic islets requires significant time and effort, and limited experiments can be performed because of their short culture life, usually no more than a week before significant reductions in functionality are evident [6]. This is equally true for primary human islets which are available for research purposes from islet transplantation programmes. These limitations have restricted investigations, such as high-resolution live imaging of islets over prolonged periods or repeated functional assessments, owing to the short culture life and insufficient techniques for culturing adherent primary islets. Other alternatives, such as β-cell lines, including MIN6 cells (mouse), EndoC-βH1 (human) or EndoC-βH5 (human), can be cultured on glass, but they are transformed β-cell lines and therefore lack the signalling input of non-β cells, including other hormone-producing cells and immune cells [7]. Although two-dimensional culture of dispersed islet cells is possible on plastic surfaces, it requires a coating with cell-derived extracellular matrices. This includes extracellular matrix secreted from human carcinoma cells or bovine corneal epithelial cell matrix, which have high batch-to-batch variability and are linked to β-cell dedifferentiation ([8-10].

In recent years, 3D culture of organ-like structures has paved the way for understanding the pathology of various diseases, aided drug discovery, enabled the identification of novel therapeutic targets, and enhanced personalised medicine. Therefore, developing novel 3D models of pancreatic islets is of high importance to aid the study of the pathophysiology of diabetes and to better understand the response to various compounds and therapeutics. Dispersion of whole islets and their subsequent reaggregation into spheroids offers one such method with the advantage of adherent long-term culture under suitable conditions and the potential for more efficient transgene delivery. A previous study attempted to use reaggregation and the subsequent formation and maintenance of islet organoids by using novel expansion and proliferation culture media. These islet clusters harboured key islet hormone-producing cell types and showed a higher degree of proliferation than native islets. However, this paradigm has limitations such as: (i) islet-derived cell clusters could only be generated from pregnant mice or wild-type rats, (ii) cluster numbers were small and inadequate for transcriptomic studies, (iii) the islet clusters were not functionally evaluated, and (iv) the presence of a resident immune population was not determined [11]. Stem cell-derived pancreatic organoids are an alternative. These cells are generated by directed differentiation of pluripotent stem cells and can be cultured over the long term. However, they also suffer from limitations, including not exhibiting all hormone-producing cell types, non-physiological insulin secretion, and a lack of a resident immune population.

Given the need for alternative methods for the long-term culture of primary islets, we aimed to refine reaggregation techniques to generate a superior adherent 3D islet culture platform using native murine or human islets as the source. These reaggregated islets are referred to as reformed islets. They can be maintained in culture for at least 4-6 weeks, making them ideal for long-term studies, including high-resolution imaging, suitable for repeated functional assessment, and studying cell-cell communication. We found the reformed islets to be architecturally, transcriptionally, and functionally highly similar to the native islets they are generated from.

## Methods

### 1.1 Materials

Hanks’ Balanced Salt Solution and Hepes buffer were obtained from Gibco, B27 (Thermo Fisher Scientific). Fluoxetine hydrochloride, collagenase type XI, histopaque-1077, culture media, and supplements were purchased from Sigma-Aldrich. The primary antibodies used in this study are summarised in Supplementary Tables 1 and 2. Recombinant murine tumour necrosis factor α (TNF-α), interferon γ (IFNγ), and interleukin-1β (IL-1β) were purchased from PeproTech EC, and the Caspase-Glo 3/7 Assay Kit was purchased from Promega.

CD8^+^ cells were isolated from the spleens of four 8-week-old diabetic female NY8.3 NOD mice using a MojoSort Mouse CD8 T-cell isolation kit (Cat# 480043, BioLegend, San Diego, CA, USA) and MojoSort buffer (Cat# 480017, BioLegend). Cell Biolabs’ CytoSelect™ cell migration assay from Cell Biolabs and EZCellTM cell migration/chemotaxis assay kits were purchased from BioVision.

### 1.2 Animal Husbandry

CD-1 male mice were purchased from Charles River and were housed under temperature-(22±2°C) and light-controlled conditions (12 h light:12 h dark cycle) with ad libitum access to drinking water and a standard rodent chow diet. Spleens for CD8+ cell isolation were obtained from four 8-week-old female diabetic NY8.3 non-obese diabetic (NOD) mice. Animal experiments were performed in accordance with the British Home Office Animal Scientific Procedures Act (1986).

### 1.3 Human Donors

Human islets were retrieved from non-diabetic, heart-beating, and brain-dead donors and isolated by cold collagenase digestion ([12, 13] with appropriate ethical approval from the King’s College London Human Islet Research Tissue Bank (KCL HI-RTB; 20/SW/0074). Islets from two donors were used in this study. Donor 1 was 41yr old female with a BMI of 24 and donor 2 was 36yr old male with a BMI of 23.

### 1.4 Islet Isolation

Mouse islets were isolated from male CD-1 mice (≥ 20 g) aged 8-12 weeks and separated from exocrine tissue by digestion with collagenase. Isolated mouse islets were cultured overnight (37°C, 95%air/5%CO2) in RPMI supplemented with 10% (v/v) foetal bovine serum (FBS), 100U/ml penicillin and 100μg/ml streptomycin, and 2mM L-glutamine.

### 1.5 Preparation of matrix-coated coverslips

Round borosilicate glass coverslips (12 mm diameter and 0.17 mm thickness) were transferred to 24-well plates and coated with either laminin from Engelbreth-Holm-Swarm murine sarcoma basement membrane at 50μg/ml or Collagen IV stock solution 1 mg/ml in Ca^2+^/Mg^2+^supplemented with Hanks Balanced Salt Solution (HBSS) overnight at 37°C. The coverslips were washed in HBSS and left to dry for 10 min before seeding.

### 1.6 Preparation of reformed islets

2000-2500 mouse or human islets were hand-picked from suspension cultures 24h after isolation and washed with PBS. Islets were then dissociated into a suspension of single cells through trypsin digestion, which was stopped by adding neurobasal media consisting of MEM supplemented with 5% (v/v) FBS, 1x B-27, 1%(v/v) penicillin and streptomycin, HEPES 10 mM, Glutamax 1x, Na-pyruvate 1mM, and 11 mM glucose. Islet cells were seeded at a density of 35,000 cells/cm^2^ on laminin-coated or collagenase-coated glass coverslips. They required 3-4 days of culture to adhere and spread on the glass surfaces, and 10-14 days to form reformed islets. These reformed islets were used for all the experiments described here. The reformed islets required a change in the media every 2 days.

### 1.7 Immunofluorescence and confocal microscopy

Immunofluorescence was used to study the cytoarchitecture of the reformed islets, as well as proliferation and cytokine-induced damage. Coverslips with 6-10 reformed islets were fixed with 4% PFA and incubated in blocking buffer (1%BSA + 10%donkey serum in 0.1% PBST) for 1h at room temperature to eliminate non-specific antibody binding. Thereafter, the reformed islets were incubated with primary antibodies against insulin, glucagon, somatostatin, Ki67, CD80, IBA1, CD8, CD3e, and CD4 overnight at 4°C and washed in PBS (Supplementary Table 1). The slides were subsequently exposed to appropriate secondary antibodies for 1h at room temperature and washed with PBS (Supplementary Table 2). DAPI Fluoromount-G was used for the mounting. Images of the reformed islets were taken with an Eclipse Ti-E Inverted A1 confocal microscope or Zeiss LSM700 Confocal Microscope and analysed using CellProfiler and ImageJ software.

### 1.8 Insulin secretion from mouse and human reformed islets

Insulin secretion from reformed mouse and human islets was measured by static incubation experiments. Reformed islets were pre-incubated for 1h in physiological salt solution (Gey & Gey, 1936) supplemented with 2mM glucose. Coverslips with 6-10 reformed islets were counted under a Nikon TMS phase-contrast microscope and incubated with appropriate volumes (100µl of buffer per reformed islet) of Gey Gey buffer supplemented with either 2mM or 20mM glucose. The supernatants were collected and stored at -20°C prior to insulin radioimmunoassay.

### 1.9 Apoptosis: Caspase-Glo 3/7 assay

Caspase-Glo 3/7 assay was used to detect apoptosis by measuring caspase 3/7 activity. Coverslips having 4-5 reformed islets and primary islets were exposed to a mix of pro-inflammatory cytokine cocktail (CK) (0.05U/µl IL-1β, 1U/µl TNF-α and 1U/µl IFN-γ) diluted in RPMI supplemented with 2% FBS and 1% Pen/strep for 24h. Caspase 3/7 activity was quantified by bioluminescence according to the manufacturer’s protocol.

### 1.10 Islet destruction: Migration assay

The Transwell cell migration assay measures the chemotactic capability of cells toward a chemo-attractant. First, we isolated islets from CD1 mice, BALB/c mice, and human cadavers. BALB/c mice were used, as these mouse strains express the MHC class H2Kd haplotype and are therefore matched to antigen-specific T cells.

The reformed islets were incubated in RPMI supplemented with 2% FBS and 1% Pen/strep for 24h. Subsequent studies using the selective serotonin reuptake inhibitor fluoxetine were chosen for migration and invasion studies, as there is evidence to support the anti-inflammatory and β-cell-protective properties of this drug [11, 14].

Simultaneously, RAW 264.7 cells were activated with LPS (200 ng/ml) and IFN-___ (2.5 ng/ml) for 24 h [15]. After washing with PBS to remove LPS and IFN-___, the RAW cells were trypsinised, pelleted, seeded (1X10^6^ cells) into inserts (upper chamber), and placed over coverslips with at least 3-4 reformed islets for an additional 24h.

CD8+ cells were isolated from the spleens of four 8-week-old diabetic female NY8.3 NOD mice. Briefly, a single-cell suspension was prepared with MojoSort buffer at 1 × 10^8^ cells/ml. 10 µl of biotinylated antibody containing antibodies against CD4, CD11b, CD11c, CD19, CD24, CD45R/B220, CD49b, CD105, I-A/I-E, TER-119/Erythroid, and TCRγδ were added. The mixture was then incubated for 15 min on ice, and 10 µL streptavidin nanobeads were added. The final volume of 120 µL of the cell and bead mixture was incubated on ice for an additional 15 min. Finally, 120 µL was diluted in in 1x MojoSort buffer (2.5 mL) and placed in a MojoSort Magnet (Cat# 480019/480020, BioLegend) for 5 min for magnetic separation. The free cells in the solution were collected as enriched CD8^+^ T cells.

The migration assay was measured using an 8µm pore transwell plate (BioVision, Inc., USA) according to the manufacturer’s protocol. Briefly, BALB/c reformed islets were incubated for 48h in 2% FBS RPMI complete medium in the lower chamber of a Transwell culture plate. Thereafter, 2X10^5^ CD8^+^ T cells were seeded into the upper chamber and placed over coverslips with reformed islets of both groups: untreated reformed islets and cytokine cocktail-exposed reformed islets. After 18 h of incubation, the cells in the lower chamber were quantified by measuring the O.D values at Ex/Em = 540/590 nm using a PHERAstar Microplate Reader.

To investigate reformed islets as a platform to perform migration studies, the effects of the drug fluoxetine on macrophage migration and reformed-islet destruction in the presence of pro-inflammatory cytokines were studied using established migration assay kits. Briefly, mouse or human reformed islets were incubated for 48h in 2% FBS RPMI complete media with or without a drug of interest for the first 24h. After the initial 24h, activated monocyte/macrophage-like RAW 264.7, were seeded into the upper chamber (2X10^5^ cells) and placed over coverslips with reformed islets. Pro-inflammatory cytokines were added for the final 24h. After 24 h of incubation, the cells in the lower chamber were quantified by measuring the O.D values at Ex/Em = 540/590 nm using a PHERAstar Microplate Reader.

### 1.11 Islet destruction: Invasion assay

A Transwell cell invasion assay was developed to measure both cell chemotaxis and invasion of islet cells through the extracellular matrix. This method is often used for studying cancer metastasis and embryonic development [16]. To this end, we studied the effects of cytokine-induced destruction of mouse and human reformed islets in the presence of CD8^+^ cells and activated monocyte/macrophage-like cells RAW 264.7 and assessed by confocal imaging [17].

For this study, we used coverslips that underwent migration assays (described above). The coverslips were fixed and immunolabelled with insulin, CD8, or CD80 antibodies and counterstained with DAPI for the identification of nuclei. Qualitative analyses using light and confocal microscopy were used to assess the appearance of the reformed islets.

### 1.12 Bulk RNA-sequencing

Total RNA was extracted from three replicates of native human islets and reformed islets from one donor, using the TRIzol method. Samples were uniquely barcoded and pooled into one sample. After sequencing, all the samples were distinguished based on their unique barcodes. The next step included In Vitro Transcription, is a linear amplification step which results in amplified RNA (αRNA). This α-RNA was fragmented and run on an Agilent TapeStation to assess RNA integrity.

Reverse transcription, in addition to a PCR reaction, amplified the α RNA, and appropriate sequencing adapters were added. During the PCR reaction, the material was again amplified, and appropriate adapters for sequencing were added to the sequences. This resulted in a DNA library (cDNA) that could be used for sequencing. The cleaned cDNA library was run on an Agilent TapeStation instrument.

Library preparation and RNA sequencing were performed by Single-Cell Discoveries using the QuantSeq 3 kit (Lexogen) and sequenced on an Illumina NextSeq 500 with a sequencing depth of 10 million reads per sample. Raw sequencing data were aligned against the human reference genome using STARsolo (2.7.10a). Read quality checks were carried out using FastQC, after which BBDuk from the BBMap (version 38.87) software was used to trim Illumina adapters and polyA tails. Reads were mapped to the hg38 reference human genome using Star Aligner.

### 1.13 Data processing

#### 1.13.1 Bulk RNA-Seq analysis

The R package DESeq2 v1.36.0 [18, 19] was used to determine differential gene expression, and DEGs with padj <0.1 were selected. Variance-stabilising transformations of raw count data were used to visualise the sample clustering. All data were subjected to global-scale normalisation and log-transformation. Linear dimensionality reduction was performed using principal component analysis on the top 500 genes from the variable-stabilising transformation dataset using the DESeq2 package. The data was also subject to shrinkage estimation of the log_2_ fold changes using apeglm.

#### 1.13.2 Bioinformatic pathway analysis

Gene Set Enrichment Analysis (GSEA) using the Hallmark, Gene Ontology biological processes, and KEGG databases was conducted with the fgsea package v1.22.0 [19]. Heatmaps were generated using normalised counts and Z-scores scaled using the Complex Heatmaps package v2.12.1 [20]. Volcano plots were generated using SRPLOT software, using a log_2_ fold-change of specific genes in the dataset. A gene was considered significantly differentially expressed if it demonstrated an adjusted p-value < 0.05 and log_2_ FC> 1.5. Sequence data are available from the GEO database under accession number GSE248341.

### 1.14 Statistical analysis

Statistical analysis and graphical representation of the data were performed using GraphPad Prism software (version 9.5.1; GraphPad Software, La Jolla, CA, USA). Statistical comparisons were performed using one-way ANOVA, followed by Šídák’s or Dunnett’s multiple comparison tests. All results are expressed as the mean ± SEM. Statistical significance was defined as a p-value < 0.05 and is denoted as stars in the graphs (* p < 0.05; ** p< 0.01; *** p< 0.001; **** p< 0.0001) in all figures, unless otherwise stated.

## Results

### 3.1 Reformed islets present organotypic cytoarchitecture akin to native primary islets

We optimised the reaggregation of islet cells by integrating a previously published protocol [10] with several enhancements and a more extensive validation process. The critical modification involved extending the maturation time after cell disaggregation and seeding from 4 days to 14 (Fig. 1). This extended period enabled the cells to re-aggregate and develop into 3D spherical structures, whereas they remained as a monolayer using the previous protocol. Furthermore, once the reformed islets reached maturity, they were successfully maintained on the platform for 42-days. This extended culture time allowed us to observe the long-term viability and functionality of the reformed islets.

**Figure 1.**
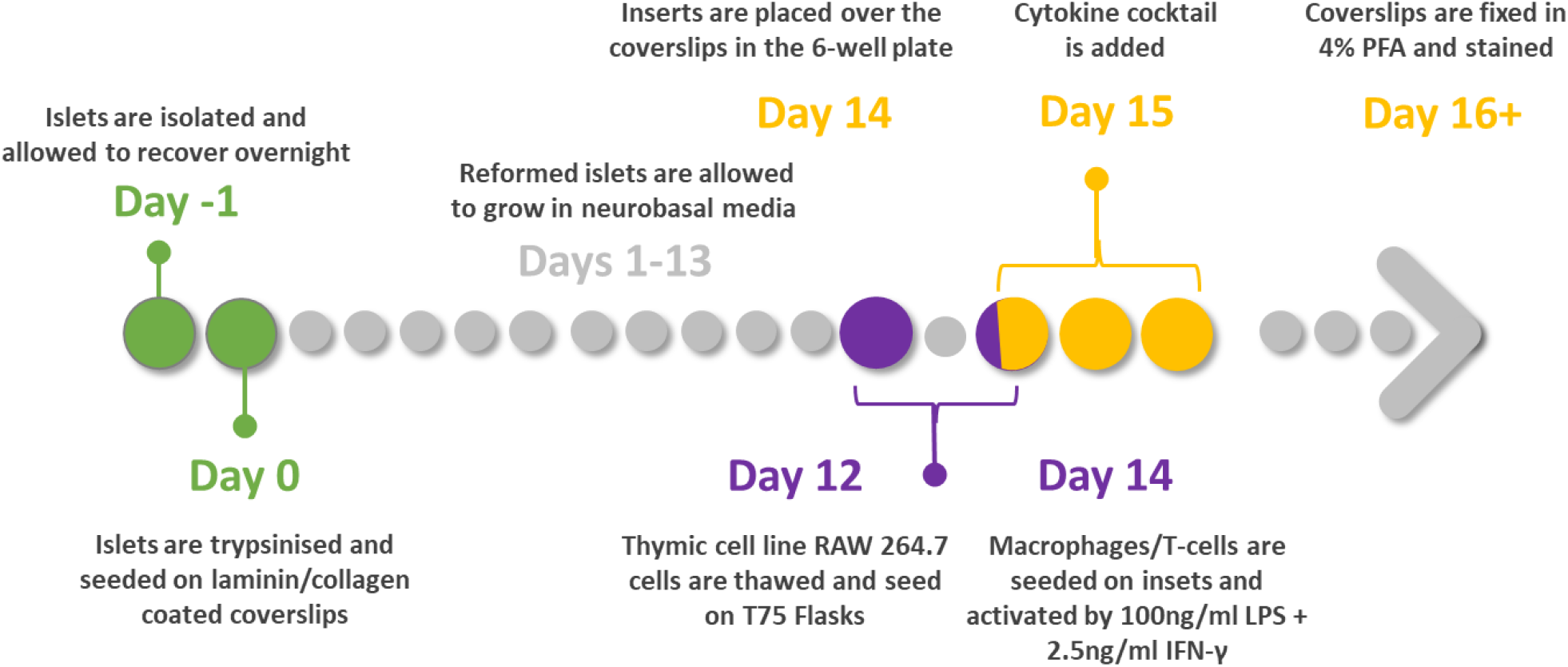
Overview of reformed protocol. Native islets isolated from mouse or human donors were allowed to recover overnight. On day 0 they were trypsinised and seeded at a density of 35,000 cells/cm^2^ on laminin-coated or collagenase-coated glass coverslips. They required 3-4 days of culture to adhere and spread on the glass surfaces, and 10-14 days to form reformed islets. Experiments were performed thereafter.

After seeding, single islet cells adhered to the coated coverslip, and within several days, these cells formed a nonconfluent but connected network. After seven days of culture, focal points within the network began to form 3-dimensional spheroid structures (Fig. S1A) that harboured multi-hormonal cells producing insulin, glucagon, and somatostatin (Fig. S1B). At 14-16 days, these spheroids were fully formed and exhibited a native islet cytoarchitecture with distinct hormone-producing populations. The size of the reformed mouse islets was comparable to that of the native mouse islets, whereas human reformed islets tended to be slightly smaller than their native counterparts (181.1±2.07 vs 126.3±18.71) (Table 1) (Fig. 2A, B).

**Table 1.**

| Charecteristics | Mouse Native Islets | Mouse Reformed Islets | Human Native Islets | Human Reformed Islets |
| --- | --- | --- | --- | --- |
| Size | 192.8±13.54 | 173.7±19.11 | 126.3±18.71 | 181.1±20.7 |
| beta cells | 76.53±2.96 | 72.64±2.54 | 65.79±9.16 | 66.38±6.39 |
| alpha cells | 15.53±3.34 | 8.951±1.57 | 26.56±2.21 | 26.73±1.52 |
| delta cells | 11.75±1.77 | 4.33±0.52 | 12.31±4.16 | 9.225±1.25 |

**Figure 2.**
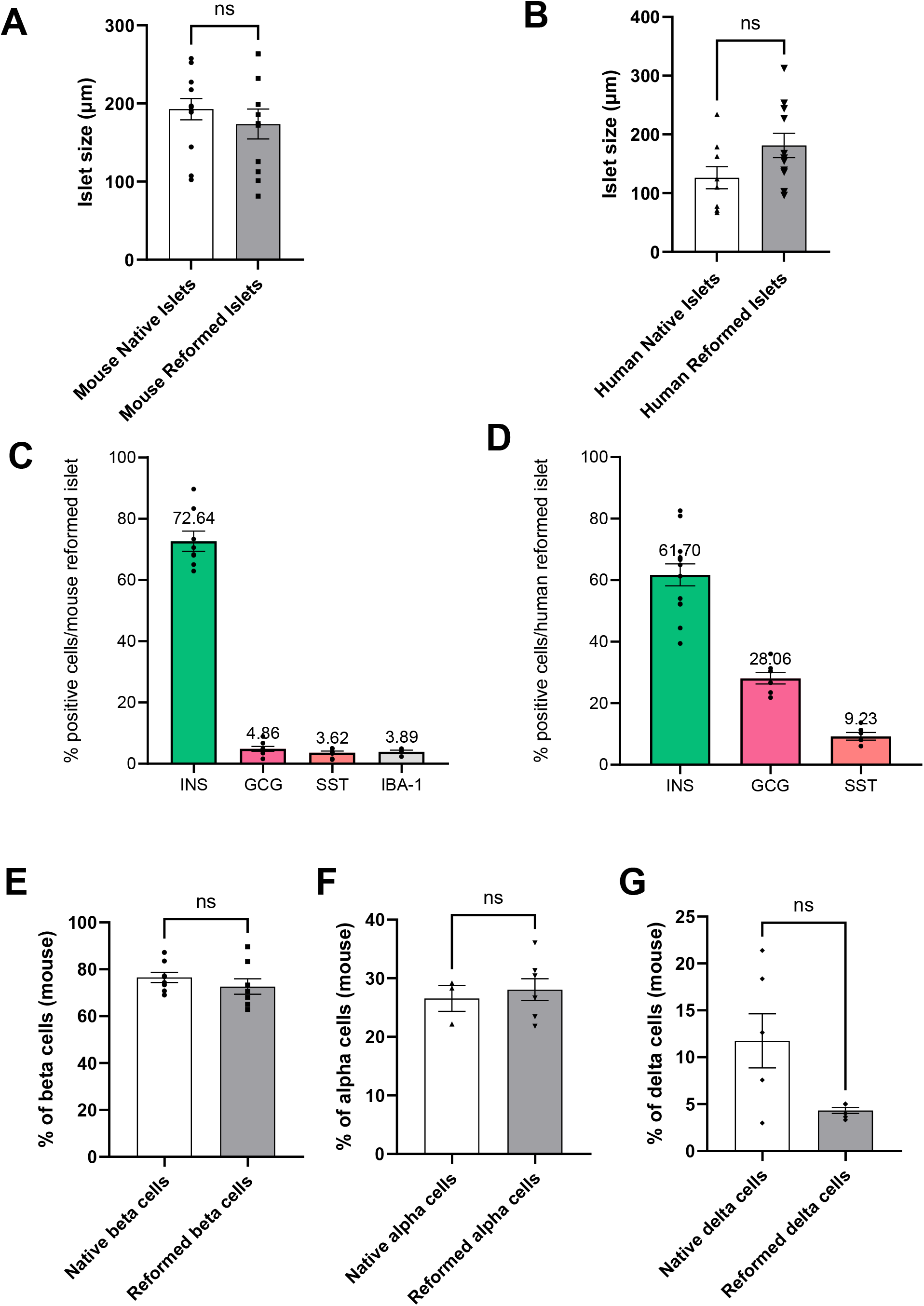
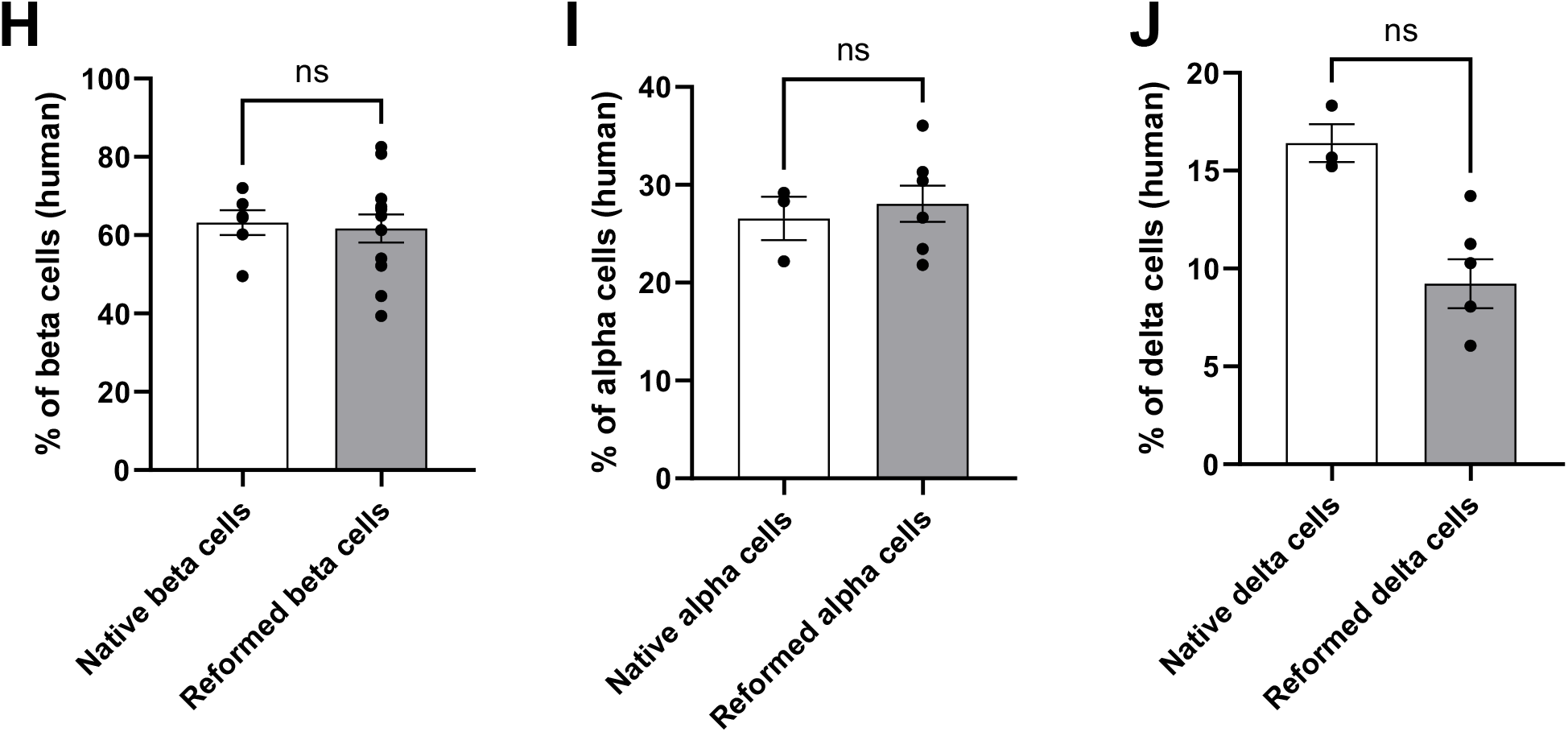
Comparative analyses of size and cell composition of native and reformed islets. The native and reformed islets were immunostained for insulin, glucagon somatostatin and IBA analysed using the ImageJ software. Sizes of native and reformed mouse (A) and human (B) islets were compared and presented in Table 1. Percentages of different cell types in mature mouse (C) and human (D) reformed islets were calculated and compared to native islets (mouse:E-G, human: H-J). Numerical data are presented as the mean ± SEM, n=4-13. Ns represents p>0.05, unpaired Student t-test or One-way ANOVA, Dunnett’s multiple comparisons test.

We evaluated the cellular composition of mouse reformed islets, with β cells comprising 72.64%, α cells constituting 4.86%, and δ cells comprising 3.62% of the total cell population (Fig. 2C). Notably, the distribution of glucagon-positive cells in the reformed islets was similar to that observed in the native islets and was localised predominantly in the periphery. Similarly, in human reformed islets, the cellular composition (β-61.70%, α-28.06%, δ-9.23%) closely resembled that of native human islets which contain 50-60% β cells, 30-50% α cells and 9-10% δ cells (Fig. 2D).

Native mouse islets consist of approximately 65–80% insulin-producing β-cells which are primarily located in the inner core and, 10–20% of the cells are glucagon-producing α cells found in the outer region of the islet. Next, we compared the cellular composition of the major cell types between native and reformed islets in both mouse and humans. In mouse, the percentages of β (*p-val*=0.48) and α cell types (*p-val*=0.41) were similar to those of reformed islets. The percentage of δ cell type was slightly lower (*p* =0.1) (Fig. 2E, F and G). In human native and reformed islets, the percentages of β-cells (*p* =0.98) and α-cells (*p* =0.99) were similar. However, there was a lower percentage of δ cells in the reformed islets (*p* =0.59) (Fig. 2H, I and J).

Similar to mouse reformed islets, the distribution of human glucagon-positive and somatostatin-positive cells mirrored that of native human islets, where they were interspersed with β-cells (Fig. 3 A, B). The hormonal cell composition and positional pattern in both mouse and human reformed islets support the notion that our modified platform is an excellent mimic of primary native islets. In addition to hormone-producing cell types, a resident immune population is present in both murine and human islets [32]. This immune population is key to normal function and accounts for approximately 2-3% of all islet cells. Macrophages (MΦ) are the predominant fraction in both mouse and human islets, but other immune cell types, such as CD4 and CD8 T cells are also present (Fig 3A, S2 C-E). Increasing evidence suggests that these immune cells play crucial roles in maintaining islet homeostasis, fine tuning islet development, and modulating insulin secretion [20, 33]. Moreover, the resident immune population has been shown to contribute to the initiation and progression of diabetes. For instance, the resident macrophage population is believed to be one of the key initiators of islet T cell infiltration, a hallmark of T1DM. The immune population in the reformed islets was verified by quantitative PCR. The transcriptional level of CD45 (Fig. S2A), and macrophages (Fig. S2B) were comparable to those of the native islets. Additionally, immunolabelling of the reformed islets revealed the presence of both resident macrophages (identified by Iba1, ionised calcium-binding adaptor molecule 1, and positivity) and resident T-cells (identified by CD8 and CD4 immunoreactivity) (Fig. 2C, and 3A-lower panel). The presence of a resident immune population is an important advantage of our platform because all other long-term culture paradigms lack these cell types.

**Figure 3.**
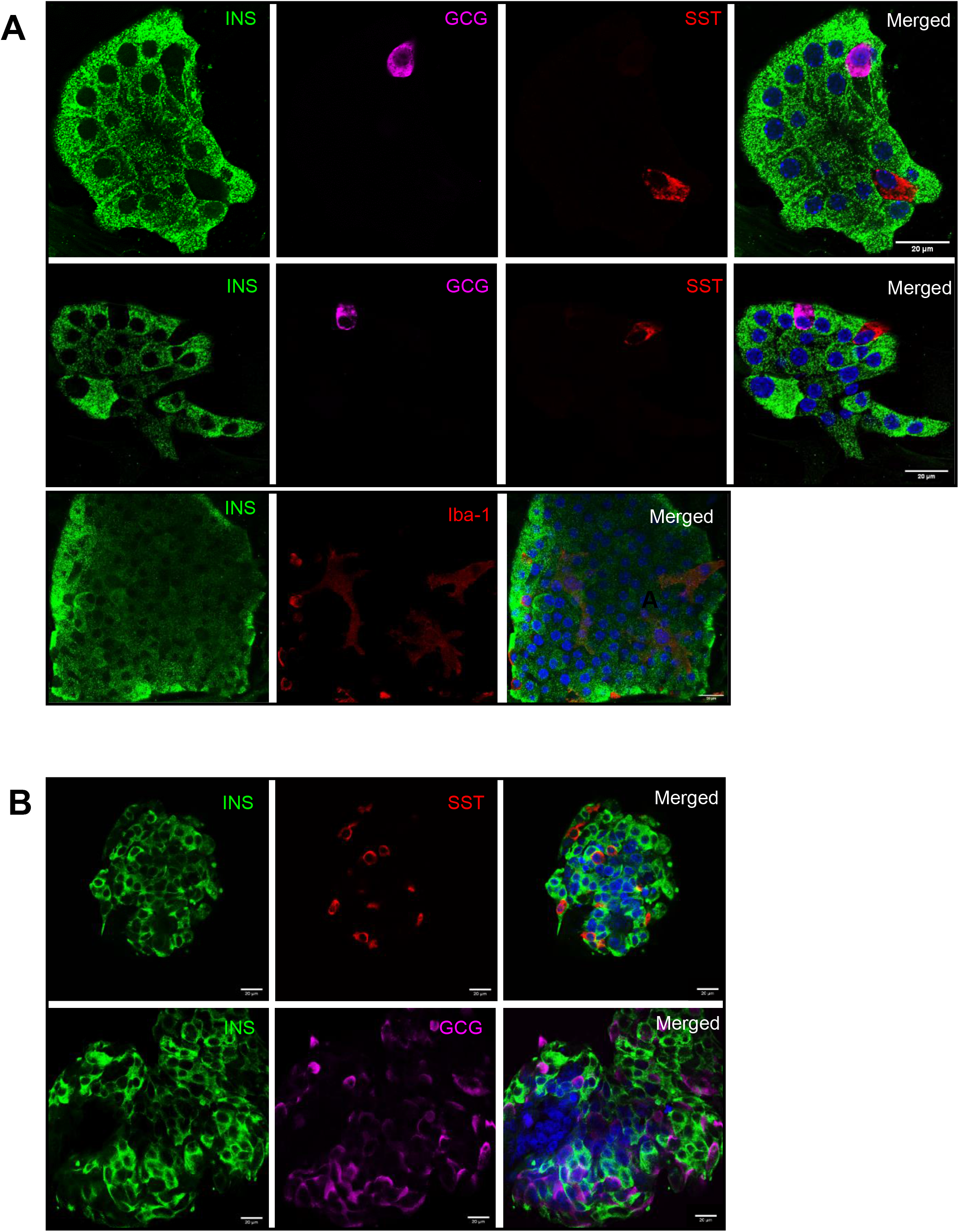
Immunofluorescence detection of islet hormones in mature mouse (A) and human (B) reformed islets formed from single islet cells 14 days after seeding. The reformed islets were immunoprobed with antibodies against insulin (INS; green), glucagon (GCG; purple), and somatostatin (SST; red). DAPI nuclear staining is shown in blue. Scale bar is 20µm. Percentage calculation of cell types in mature mouse (C) and human (D) reformed islets was performed using the ImageJ software. Numerical data are presented as the mean ± SEM, n=6-13.

### 3.2 Reformed islets are able to proliferate when exposed to a GLP-1 receptor agonist

Mature β-cells within primary islets typically remain in a quiescent state. However, one potential approach to increase functional β-cell mass to treat insulin-dependent diabetes is to induce proliferation of the residual β-cell population. We investigated whether reformed islets retained the ability to proliferate when treated with the GLP-1 receptor agonist exendin-4, which is known to enhance β-cell proliferation [21]. Exposure to exendin-4 significantly increased islet Ki67 immunolabelling (*p* <0.05), which is a marker of cell proliferation (Fig 4A, B). This indicates that reformed islets exhibit a capacity for proliferation, highlighting their utility in elucidating ways to promote β-cell mass expansion.

**Figure 4.**
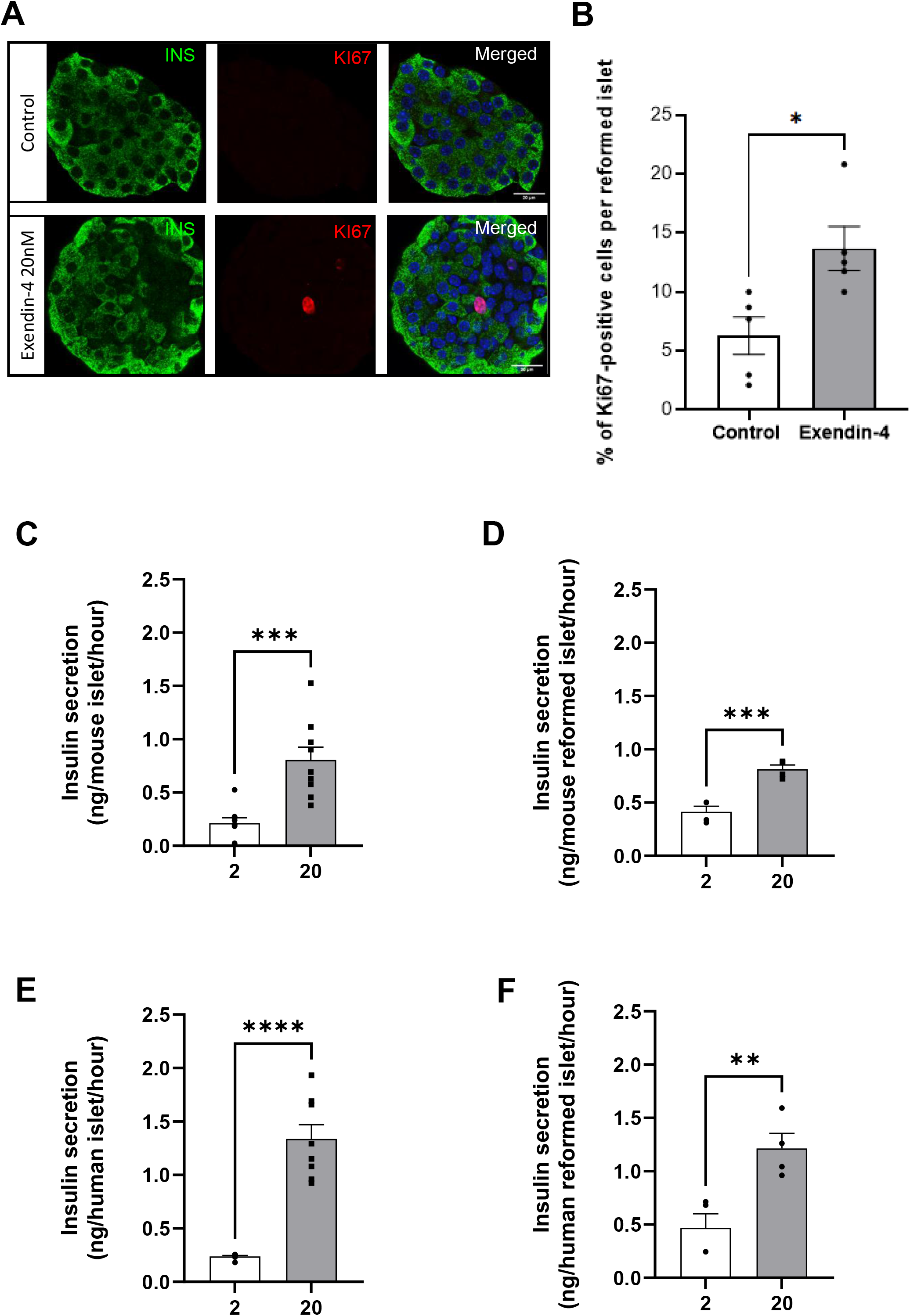
Effects of exendin-4 on proliferation of mouse reformed islets. (A)The reformed islets were exposed to exendin-4 for 30h and immunoprobed with antibodies against insulin (INS; green) and Ki67 (red). DAPI is shown in blue in merged images. Scale bar is 20µm. Percentage calculation of Ki67^+ve^ cells within the reformed islets was performed using the ImageJ software (B). Data are expressed as mean ± SEM, n=5 observations from two separate experiments. *p<0.05 relative to the control samples, One-way ANOVA, Dunnett’s multiple comparisons test. Insulin secretory response of primary islets (mouse – C, human – E) and reformed islets (mouse – D, human –F) to supramaximal glucose concentration (20mM). In static incubation experiments, islets or reformed islets were pre-incubated for 2h in Gey and Gey buffer supplemented with 2mM glucose and then incubated for 1h in the presence of either 2mM or 20mM glucose. All data are shown as the mean ± SEM, n=4-8 observations. *p<0.05; **p<0.01; ***p<0.001; ****p<0.0001 relative to the control samples at 2mM glucose, unpaired Student t-test.

### 3.3 Functional insulin secretion machinery in reformed islets

The dissociation of whole islets into single cells leads to the loss of connection which can cause dysregulated insulin secretion [22]. Insulin secretion in response to glucose in the islets of Langerhans is a tightly regulated process that involves multiple cell types and signalling pathways. In response to glucose, mature primary islets display a biphasic insulin secretory response [23].

Insulin secretion was quantified in static incubation experiments using primary and reformed islets. Acute exposure (1h) to supramaximal glucose levels (20 mM) resulted in significantly increased levels of secreted insulin compared to 2 mM glucose (Fig 4 C, D) in both native and reformed mouse islets. Similarly, the concentrations of secreted insulin were comparable between the native and reformed human islets (Fig 4 E, F). This suggests that the reformed islets are glucose sensing and capable of nutrient-driven insulin secretion.

### 3.4 Transcriptional Analysis of reformed islets

Human pancreatic islets were obtained from a 36-year-old male cadaver with a BMI of 23 (> 85% purity and >60% viability). Native islets were dissociated and seeded onto collagen-coated coverslips. After 15 days of culture, native islets and reformed islet samples were processed for paired bulk RNA-Seq analysis (Single-cell Discoveries Ltd.). More than 70% of raw reads were mapped to the reference genome. Globally, 1800 genes were differentially expressed with an adjusted p value < 0.05 when a cut-off value of > ±2 was applied to the log_2_ fold change (FC) (Fig.5A). Hierarchical clustering of the top 30 genes demonstrated significant differences between native and reformed islets (Fig. S3A). To further analyse these data, Kyoto Encyclopedia of Genes and Genomes (KEGG) enrichment analysis (Fig.5 B, C) was performed. Overexpression RNA-Seq analysis using significant log_2_FC discards a large proportion of the dataset that may be of biological relevance. Therefore, using the Molecular Signature Database collection of hallmark genes, we performed Gene Set Enrichment Analysis (GSEA)[24, 25] on our dataset as a complementary method. The hallmark gene dataset is a compilation of various published well-defined biological processes or states that display coherent expression. GSEA uses only p-values, thereby reducing the variation and redundancy of gene expression across various pathways.

**Figure 5.**
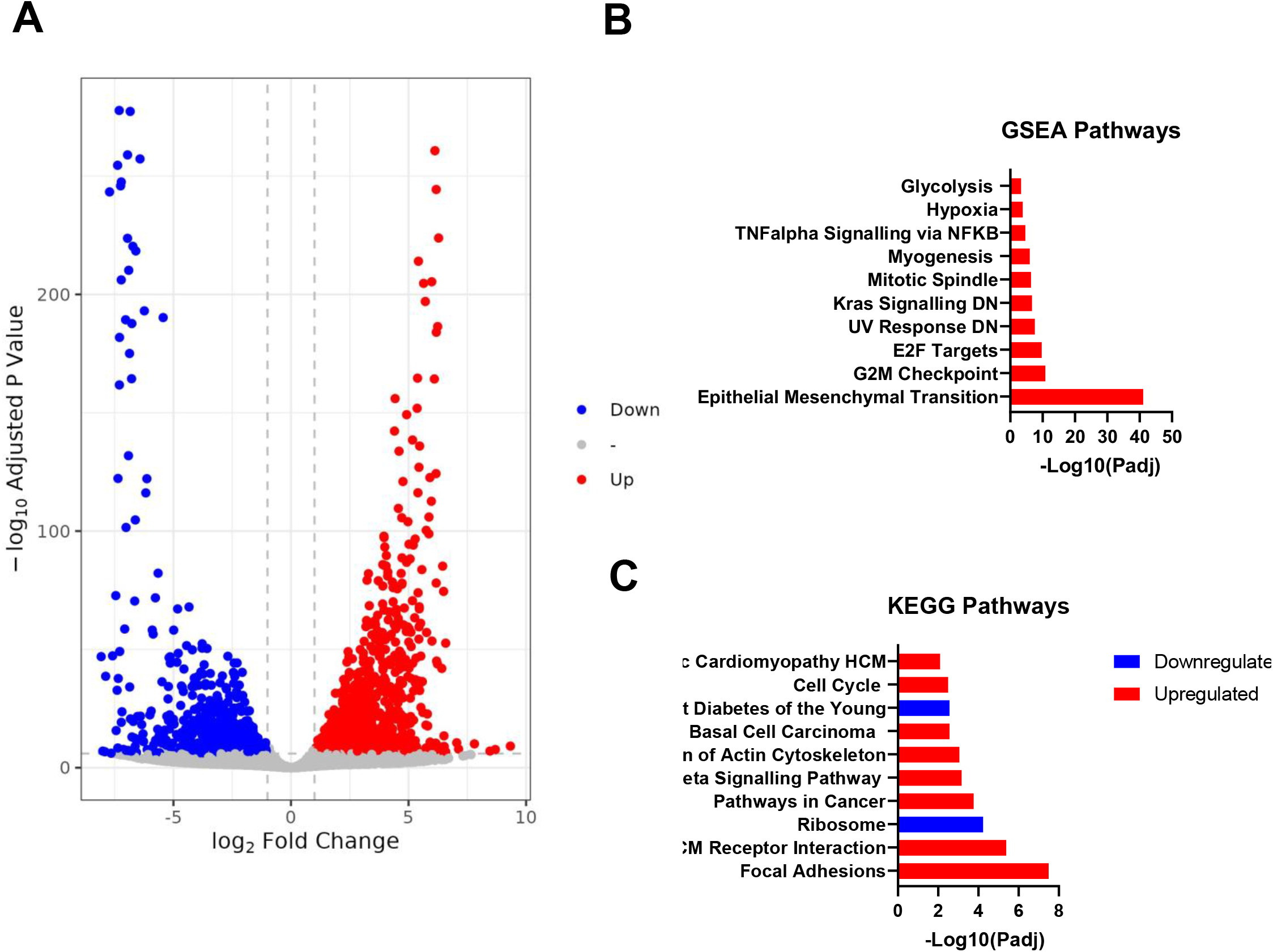
Transcriptomic profiling of reformed human islets. (A) Volcano plot of all genes significantly upregulated or downregulated in reformed islets (log2-fold-change >2, benjamini-hochberg corrected p-value threshold = 0.1). (B) Gene set enrichment analysis (GSEA) of RNA-seq data performed on reformed islets illustrates enriched pathways. (F) KEGG pathway analysis was performed for functional enrichment of genes in reformed islets.

The enriched upregulated pathways included epithelial-mesenchymal transition (EMT) (*VACN, TGFB1, THBS2, LOXL2, ACTA2, ADAM12, SGCG, POSTN, COL1A1, CDH2* etc.) (Fig. S3B), focal adhesion (*COL5A, PDGFRA, THBS2, COL1A2, FLNC, COL5A1, ITGA11, COL1A1, COMP*, etc.) (Fig. S3 C), and skeletal system development (*PDGRFA, HAND2, HOXA1, HOXA3, HOXA4, HOXA5, SULF1, HOXC6, HOXC5, COL1A1*, etc.) (Fig. S3 D). As expected, there were overlaps between genes with enriched pathways. The upregulation of transcripts encoding EMT genes, focal adhesions, and skeletal muscle development suggests that the EMT process is ongoing in cultured reformed islets. EMT has been found to be present in human epithelial cell cultures [13], and a recent study established that EMT occurs when human islets are allowed to expand in culture. These mesenchymal cells are highly proliferative and maintain the ability to differentiate into hormone-producing cells [26].

The downregulated differentially expressed genes (DEGs) were enriched in KEGG pathways related to ribosome biogenesis (*RPL11, RPL14, RPS6, RPL12, RPL21, RPL13, RPS15, RPS9, RPL3, RPS4X* etc.) (Fig. S3E), and Maturity Onset of Diabetes in the Young (MODY) (*NR5A, NKX6.1, GCK, INS, IAPP, ONECUT1, FOXAA3, FOXA2, HNF4A*, etc.) (Fig. S3F). Downregulation of ribosomal RNA (rRNA) transcription often occurs in response to intra- and extracellular stimuli in order to maintain homeostasis. Downregulation of rRNA transcription has been associated with cellular processes such as cell cycle arrest and reduced cell growth, which may positively regulate cell differentiation [27]. Genes within the MODY pathway have been shown to be positively correlated with insulin secretion, and a decrease in gene expression has been associated with β-cells being unable to maintain their mature phenotype [28, 29]. The expression of islet hormones, such as INS, GCG, SST, PYY, GHRL, and NPY, was generally mildly downregulated in reformed islets compared to their native source (Fig. S3G) despite the proportions of cell numbers being similar when assessed by immunostaining. Additionally, transcription factors related to the development of endocrine cells, such as RFX3, NKX6.1, FOXA2, and GLIS3, were downregulated (Fig. S3H). Collectively, the transcriptional landscape of the reformed islets suggests that they have a high degree of similarity to the primary islets they are derived from but exhibit a partially dedifferentiated and relatively immature state. These features may aid long-term culture stability.

### 3.5 Reformed islets response to cytokine-induced damage mimics that of primary islets

Inflammatory responses and the presence of cytokines secreted by immune and non-immune cell types are central to both types of diabetes [30, 31]. While T2DM is associated with insulin resistance and β-cell dysfunction, recent studies in mouse models have linked an increase in circulating inflammatory factors, such as chemokines and cytokines, to islet inflammation and insulitis [32]. This is mainly due to the increased number of intra-islet macrophages that secrete cytokines such as IL-1β. Three cytokines have been implicated in this process: IFN-γ, produced by CD4+ and CD8+ T cells, and TNF-α and IL-1β, produced by islet-infiltrating dendritic cells and macrophages [33]. The presence of this ‘cytokine storm’ orchestrates inflammation of islets with subsequent β-cell death in T1DM, and enhancement of insulin resistance with impairment of glucose homeostasis in T2DM.

We modelled this ‘cytokine storm’ by addition of a cocktail of pro-apoptotic cytokines (IL-1β, TNF-α and IFN-γ), and quantified caspase 3/7 activation in our reformed islets. Our results illustrated that cytokine-induced damage in mouse reformed islets (Fig. 6A) closely resembled that observed in primary mouse islets (Fig. 6B). We further qualitatively evaluated this cytotoxic damage by immunolabelling untreated and cytokine-treated reformed islets stained with antibodies directed against insulin and cleaved caspase-3, a surrogate marker of apoptosis. We observed that the untreated mouse reformed islets mostly maintained a round shape with clearly defined margins, in contrast to the cytokine-treated reformed islets that exhibited a loss of architecture with increased granularity, indicating islet destruction (Fig. 6C).

**Figure 6.**
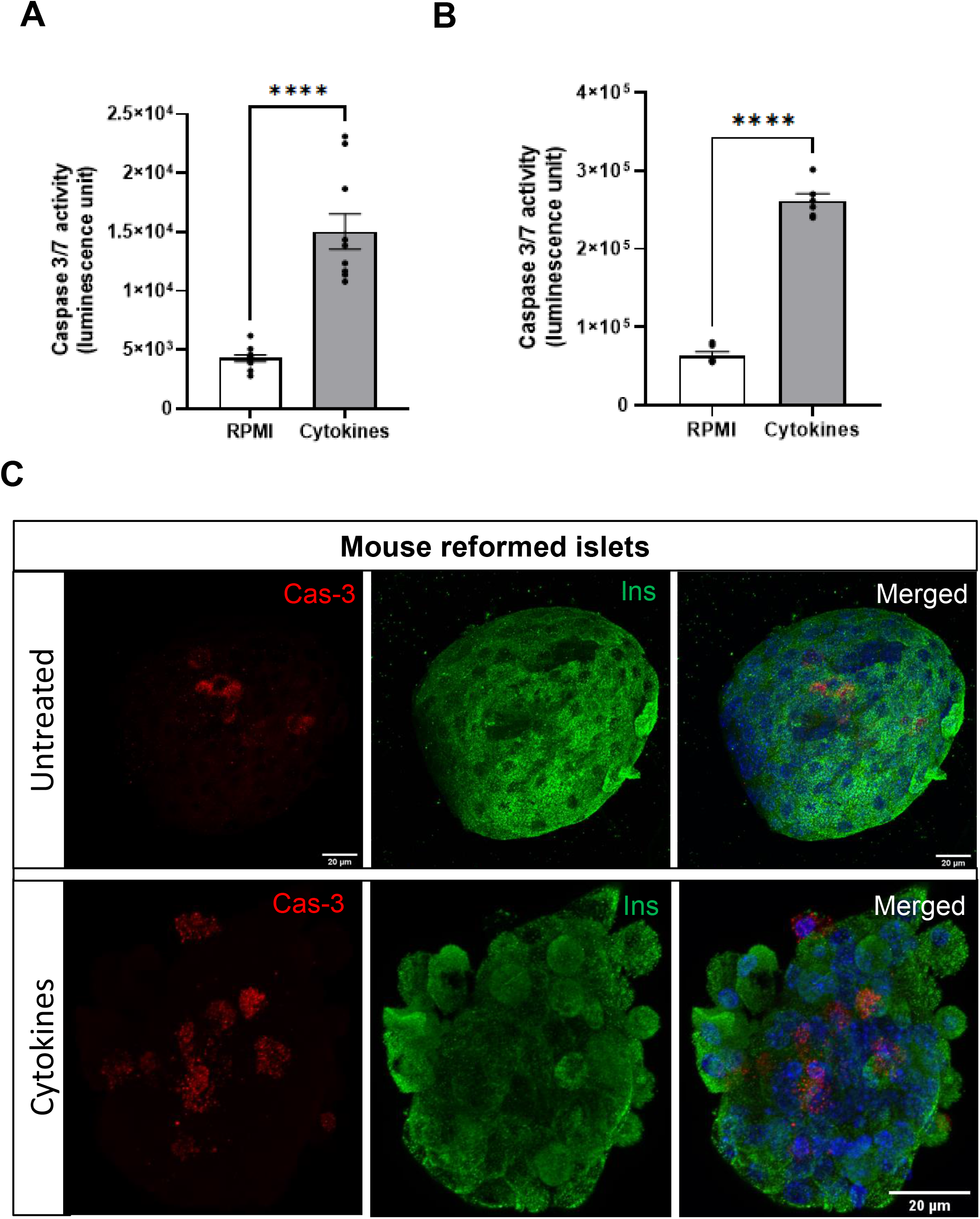
The responses of mice and human reformed islets to pro-inflammatory cytokines. Reformed islets were incubated with a mix of pro-inflammatory cytokines (IL-1β, TNF-α, IFN-γ) for 24h. Apoptosis of mouse (A) and primary islets (B) was measured by quantifying caspase 3/7 activities using the Caspase-Glo 3/7 assay following a 24h exposure to cytokines. In parallel, reformed islets were immunoprobed with antibodies against insulin (INS; green) and cleaved caspase 3 (CAS-3; red) (A,). DAPI is shown in blue, and the scale bar is 20µm. Data are shown as mean ± SEM; n=8 observations representative of three experiments. ****p<0.0001 versus control; One-way ANOVA, Dunnett’s multiple comparisons test.

### 3.6 Reformed islets are a qualitative and quantitative platform to study immune invasion and migration

The development of insulitis is a common feature in both T1DM and T2DM. Although their aetiologies differ, the invasion and accumulation of macrophages and T cells in the islets display a pro-inflammatory phenotype [43, 44] that is common to both inflammatory processes.

The progression of T1DM involves immune cell infiltration of islets, specifically CD8+ and CD4+ T cells. The exact mechanism remains unclear; however, resident antigen-presenting cells (APCs), such as dendritic cells and macrophages, detect β-cell proteins as ‘non-self’ (MHC class I molecules on β-cells), causing the processing of β-cell-specific antigens and their presentation to CD4+ T lymphocytes via the MHC class II complex [34, 35]. CD4+ T helper cells become activated, allowing them to further activate B lymphocytes and trigger autoantibody production [36]. Additionally, CD8+ cytotoxic T cells are activated through β cell antigen presentation on MHC class I, triggering the release of cytokines and resulting in β cell death [37].

The disease progression of T2DM is associated with the development of insulin resistance and β-cell dysfunction. Interestingly, recent studies in mouse models have found that the number of intra-islet macrophages increases significantly with an increase in circulating inflammatory factors, such as chemokines and cytokines, leading to islet inflammation and insulitis [32].

Using our reformed islets, we modelled immune migration and invasion in both types of diabetes by co-culturing them with either T cells or macrophages in the presence or absence of a cytokine cocktail to drive a pro-inflammatory environment (Fig. 7A).

**Figure 7.**
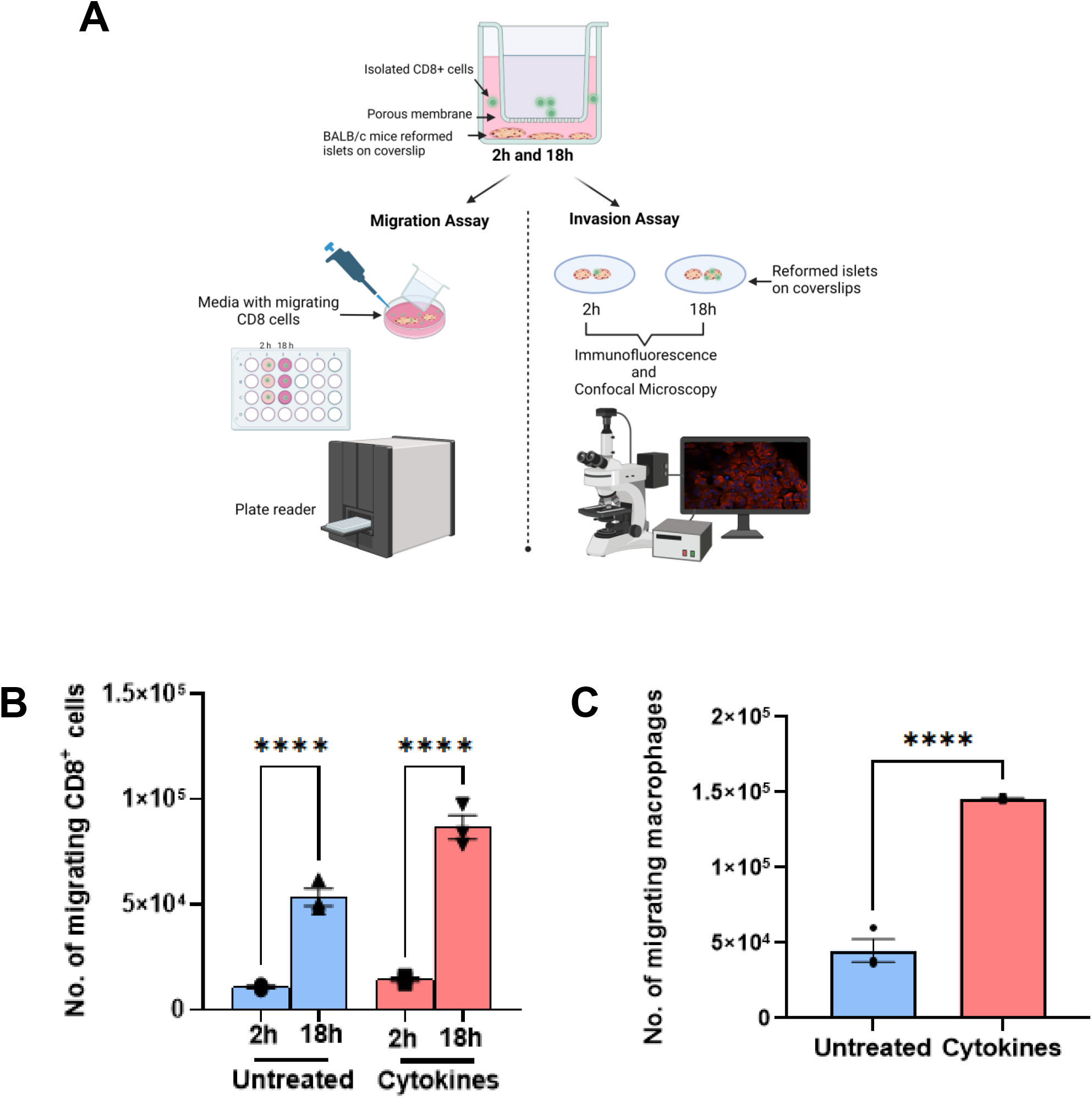
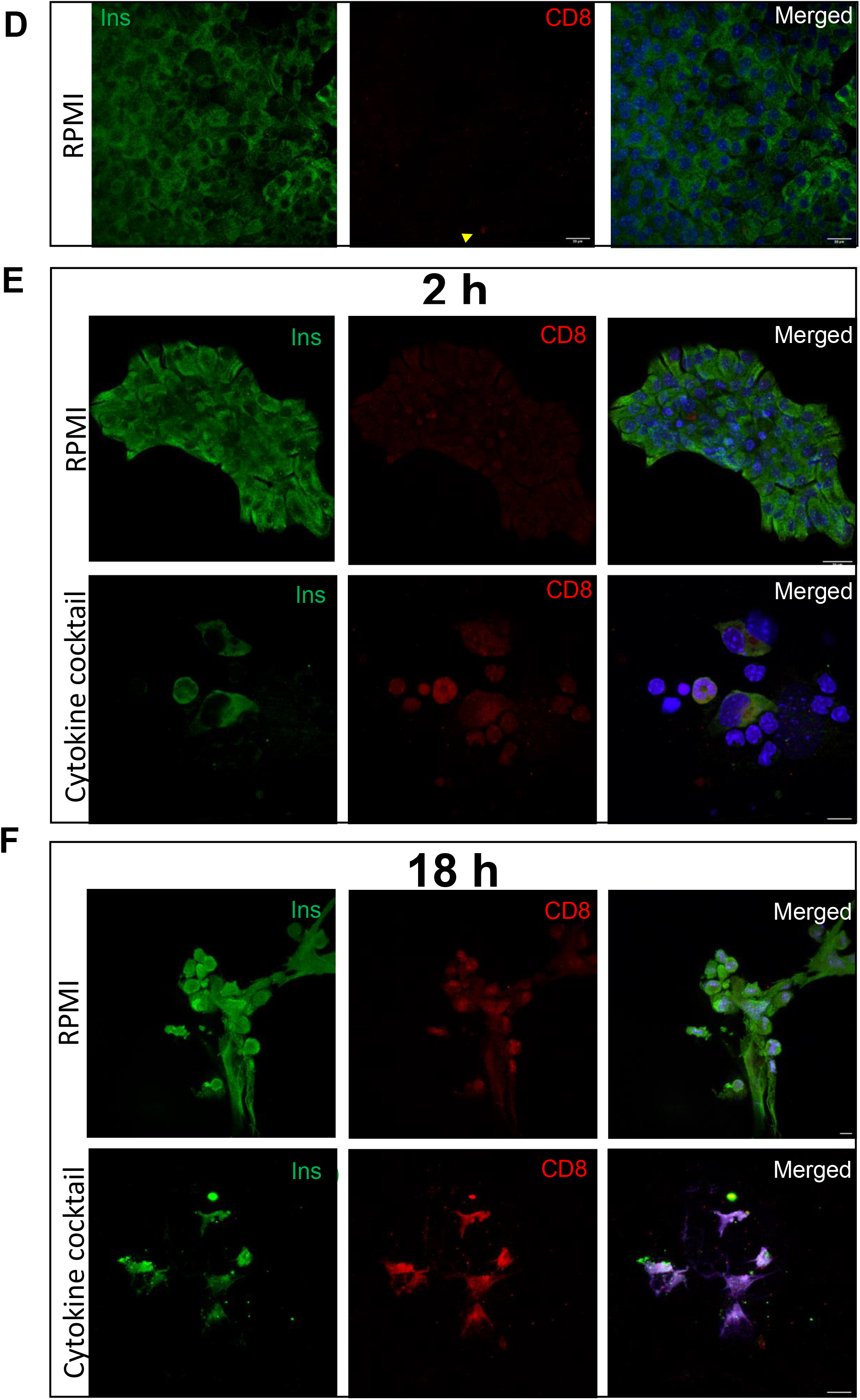
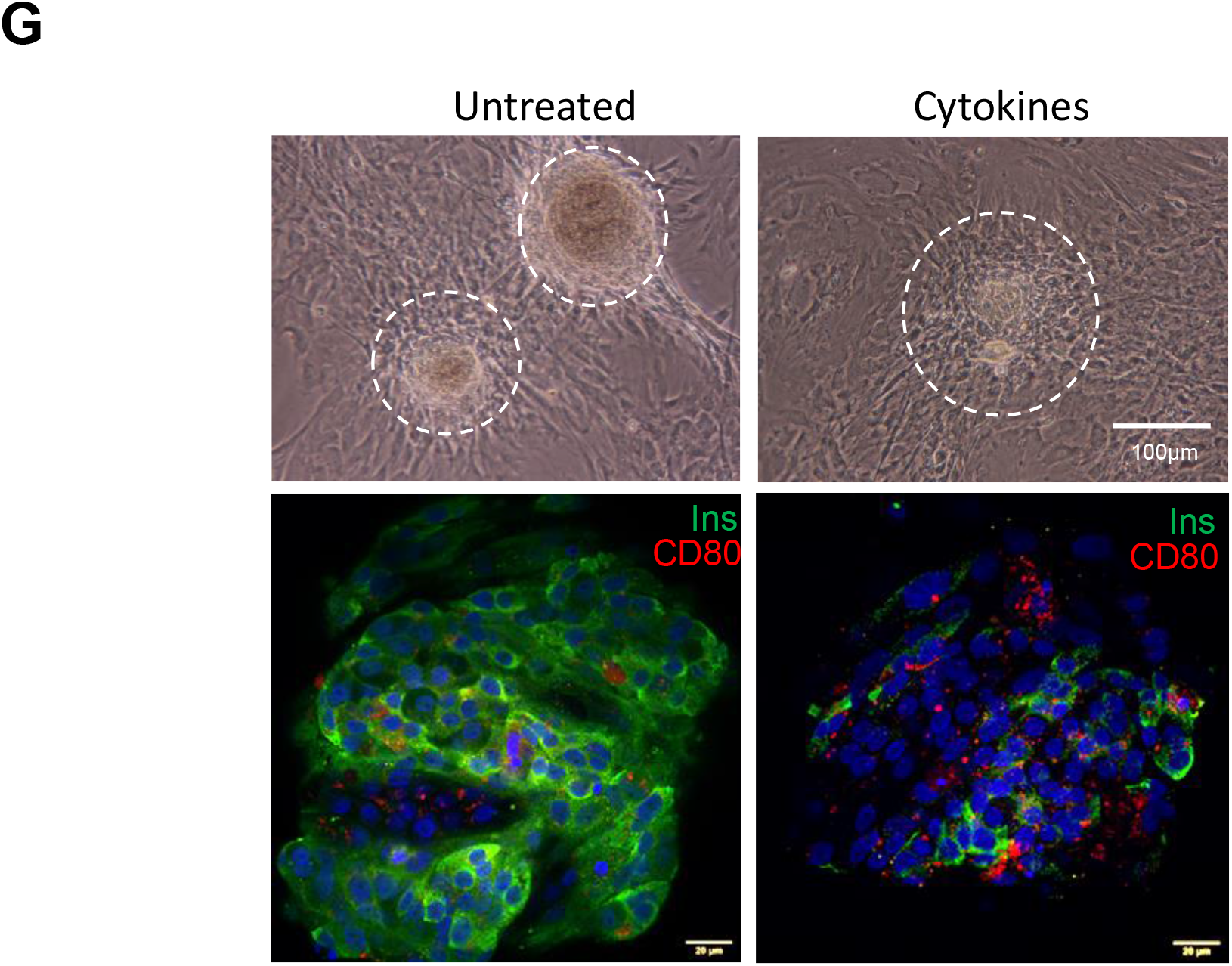
Illustration of migration and invasion assays. Schematic diagram of *ex vivo* migration and invasion assays of CD8+ T cell and activated macrophages (RAW 264.7 cells) (A). Bar graph shows mean total percentage of CD8+ T cells migrating when cocultured with haplotype matched BALB/c reformed islets (B). Bar graph shows mean total percentage activated RAW 264.7 macrophage-like cells co-cultured with reformed CD1 mouse islets and number of migrating cells☐±☐SEM. for three independent migration experiments (C). One-way ANOVA or unpaired Student t-test were performed to assess significance. ****p☐<☐0.0001. In invasion experiments, the reformed islets were immunolabelled with antibodies against insulin (Ins; green), CD8 (red) and a nuclear stain DAPI or investigated using light microscopy (7 D-G). Cytokine-induced destruction of reformed islets was investigated at time points 2h (E) and 18h (F) in comparison to non-cytokine treated (control; RPMI) reformed islets.

#### 3.6.1 Migration assay

In native islets with insulitis, CD8^+^ cells are the most abundant invading cell type and are reactive against β cell antigen-presenting cells in T1D. To mimic *in vivo* infiltration by autoreactive CD8 cells in islets, a transwell migration assay was performed to detect the migratory ability of CD8^+^ cells. Reformed islets were derived from BALB/c mice, and CD8 cells (haplotype matched) were harvested from the spleens of non-obese diabetic (NOD) mice, seeded onto the membrane of the transwell insert, and placed over coverslips with adherent reformed islets. As expected, haplotyped matched T-cells migrated to reformed islets, and this process was significantly increased during inflammation after 18h (Fig. 7B).

An increase in the number of islet-resident macrophages is prevalent in both types of diabetes [38, 39]. To model this, we co-cultured reformed islets with RAW264.7 cells a murine macrophage-like cell line. Under inflammatory conditions, RAW cell migration was significantly increased (Fig. 7C).

#### 3.6.2 Invasion assay

Next, we quantified cell migration using an invasion assay. Reformed islets were co-cultured with T cells or activated RAW cells at baseline and during the inflammatory stimulus. Invasion was imaged by immunostaining of immune cells (CD8 or CD80 antibodies) and β-cells (insulin antibody) in combination with confocal microscopy. Reformed islets derived from BALB/c mice were co-cultured with CD8^+^ T cells and qualitatively assessed at 2 and 18 h. After 2 h, we observed that reformed islets exposed to cytokines showed destruction of their architecture and infiltration of CD8^+^ cells compared with reformed islets not exposed to CD8^+^ cells (Fig. 7E). After 18 h, both cytokine-treated and untreated reformed islets were completely invaded by CD8 + cells, with complete destruction of the islets (Fig. 7F. To study macrophage invasion, reformed islets were exposed to an inflammatory stimulus and the invasive capability of LPS and IFN-γ activated RAW 264.7 cells was assessed. The number of infiltrating RAW 264.7 cells were qualitatively higher in the presence of cytokines than in untreated controls (Fig.7G).

#### 3.6.3 Islet protective agent can cloak islets from immune cell infiltration

Finally, as a proof of concept, we sought to validate the use of our platform to identify compounds that could prevent islet invasion and protect islets from damage. To this end, we assessed the islet-protective effects of fluoxetine. Fluoxetine is a commercially available anti-depressant drug that has been shown to significantly increase β-cell proliferation and protect islet cells from cytokine-induced apoptosis [14].To replicate the protective effect of fluoxetine, reformed islets were exposed to a cytokine cocktail in the presence or absence of a therapeutic concentration of fluoxetine (1 µM) for 24h while activated macrophages (RAW 264.7) were added to the inserts of the Transwell plate and cultured for a further 24h. Quantitative analysis indicated a significant reduction in the number of migrating activated RAW cells when reformed islets were cultured under inflammatory conditions and treated with fluoxetine (Fig 8A). Additional qualitative analysis by confocal microscopy illustrated the reduced migration and islet protection (Fig 8 B, C).

**Figure 8.**
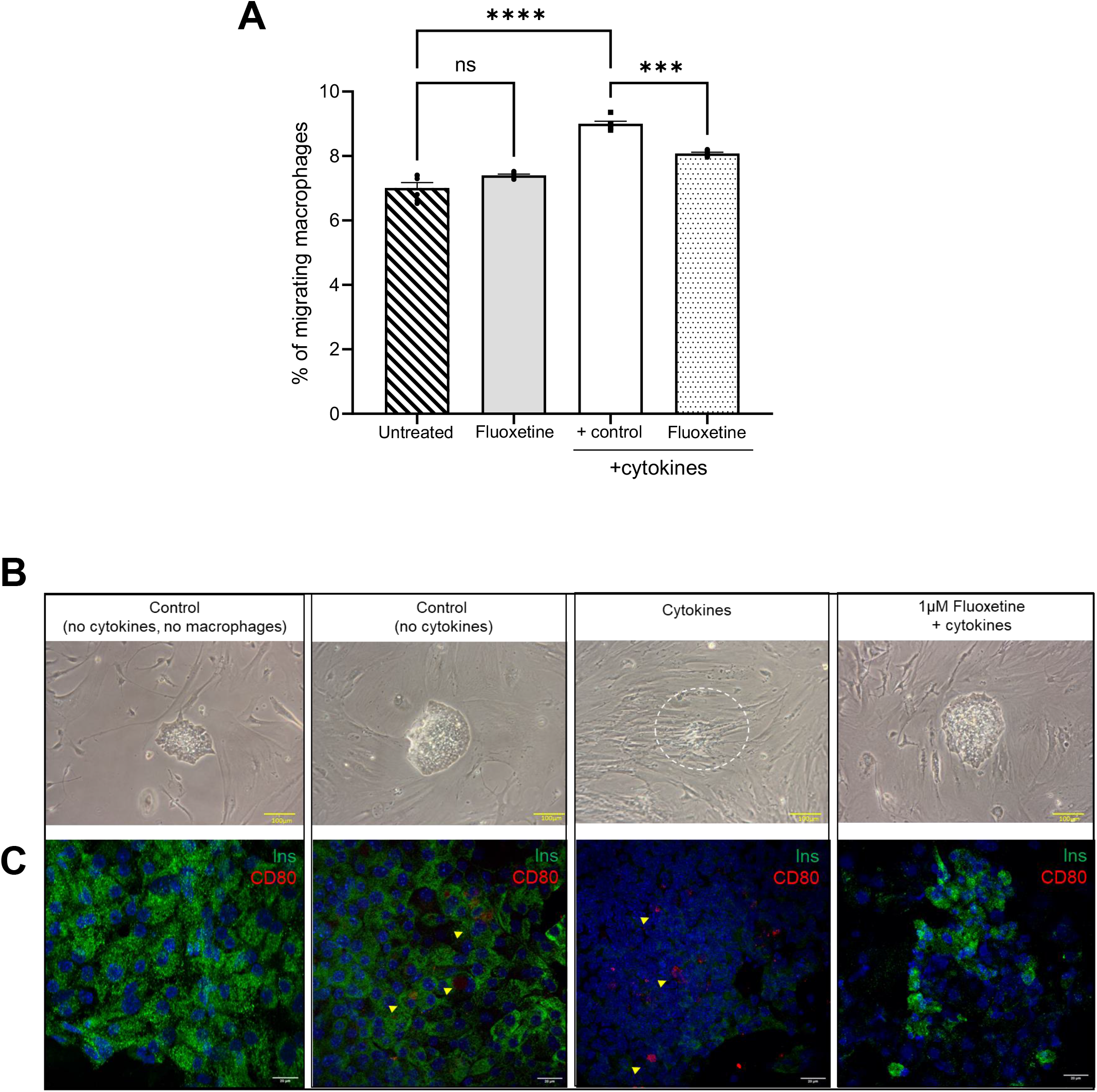
Invasion assay illustrating the effects of 1µM fluoxetine, on reformed mouse islet infiltration by RAW 264.7 macrophage cells. The reformed islets were incubated with 1µM fluoxetine for 48h and cytokines and LPS-activated macrophages were added during the last 24h of culture. Reformed islets were fixed in 4% PFA prior to imaging and fluorescence measurements were obtained (A). The reformed islets were visualised using light microscopy and immunoprobed with antibodies directed against CD80 (red), and DAPI was stained blue in parallel. Scale bars represent 100µm for light microscopy and 20µm for fluorescence microscopy. The migratory properties of RAW 264.7 cells were additionally quantified by Cell Biolabs’ CytoSelect™ cell migration and EZCellTM cell migration/chemotaxis assays (Fig.8A). Data are expressed as mean ± SEM (n = 3 observations). ***p<0.0006; ****p<0.001 versus cytokine control, One-way ANOVA, Dunnett’s multiple comparisons test.

Collectively, we show that reformed islets can be modelled to replicate the *in vivo* pathogenesis common to both T1DM and T2DM. Moreover, interrogation of reformed islets in a T2DM setting in the presence of an islet protective compound, fluoxetine, shows that this platform is ideal for exploring the function of drugs designed to protect islets from damage.

## Discussion

This study reports the development of a long-term biomimetic method to model native mouse and human islets *ex vivo*, while maintaining long-term islet functionality. We performed transcriptome profiling to evaluate how closely our model recapitulates native islets and glucose-stimulated insulin secretion, to assess functionality and evaluate islet morphology in terms of islet cell-type distribution and size. We further investigated the suitability of this platform for studying beta cell proliferation and immune cell-islet cell interactions by performing invasion and migration assays.

One of the major results of this study highlighted that reformed islets mimic primary islets and can survive for long periods in culture. In our protocol, disassociated primary islet cells are reaggregated and, over time, form 3D structures which mimic their primary counterparts.

This is in contrast to previously reported protocols, in which reaggregated islet cells remain in a monolayer [9]. Both mouse and human reformed islets harbour the major islet hormone cell types arranged architecturally, and in proportions similar to primary islets [40]. The functionality of the reformed islets was assessed using insulin secretion, beta-cell proliferation, and cytokine-induced apoptosis assays. Our results indicate that glucose-stimulated insulin secretion is comparable to that observed in primary mouse and human islets [41]. In addition, we observed that reformed-islet cells could be stimulated to proliferate in the presence of exendin-4 and that cytokine-induced damage could easily be measured using a caspase 3/7 assay [42-44]. Our findings illustrate that reformed islets are an anatomical and functional alternative to native human and mouse islets, but which can be cultured for longer periods (4-6 weeks).

To understand how closely our reformed islets modelled native islets, we performed bulk RNA sequencing and transcriptomic analysis of freshly isolated native human islets and the reformed islets on day 16. Global expression profiling revealed a number of differentially expressed genes between native and reformed islets. However, functional clustering and pathway analyses of differentially expressed genes indicated an association with Epithelial-mesenchymal Transition (EMT) hallmark genes in the reformed islets. These EMT-enriched genes have stem cell-like properties and may generate endocrine islets via committed precursor cells [45, 46]. We postulate that dissociation of native islets results in dedifferentiation and EMT activation of cells, which then redifferentiation over-time following aggregation giving rise to all endocrine cell types. It is plausible that dedifferentiation and reaggregation enables islets to withstand long-term culture conditions.

One of the key advantages of our model is the presence of a resident immune population similar to that of native islets. Various studies have highlighted the role of resident immune cells, particularly macrophages, in islet development, beta cell homeostasis, and regeneration, as well as in the pathology of diabetes [47, 48]. Indeed, the onset of diabetes, specifically T1D, results from macrophage activation which drives the recruitment of CD8+ and CD4+ T-cells to the islet, ultimately resulting in the destruction of beta-cells [49]. The lack of *in vitro* models where resident immune cells can be studied has hampered our understanding of this disease process. To our knowledge, this model is the first in which most immune cell types can be found.

Using reformed islets, we were able to recapitulate features of the immune infiltration observed in T1D mechanisms by coculturing reformed islets generated from BALB/c mice and splenic CD8^+^ T cells from haplotype-matched NOD mice. We speculate that this platform can be extended to human reformed islets by co-culturing haplotype-matched reformed human islets with human T-cell clones. Furthermore, we could also model T2D. T2D patients have an elevated number of macrophages infiltrating their islets [50, 51]. We modelled this phenotype by coculturing mouse reformed islets with macrophage-like RAW 264.7 and observed complete infiltration and destruction of islets in the presence of cytokines.

Fluoxetine, an approved antidepressant drug has recently been shown to increase insulin secretion in response to glucose and to increase beta cell mass [14, 52, 53]. As a proof of concept, we chose this drug to validate our human reformed islet model and qualitatively and quantitatively assess Fluoxetine’s ability to protect islets against cytotoxic insults. As expected, the migration and invasion assays showed that fluoxetine reduced macrophage migration and apoptosis. Thus, our study demonstrated that reformed islets can be used to identify new targets or could be employed as a screening platform for drug repurposing or repositioning once optimised in a multi-well format [54].

Long-term imaging of pancreatic islets is challenging, and the gold standard method for measuring β-cell mass is to evaluate the pathological staining of fixed pancreatic tissue sections. However, this does not encompass the dynamic changes that the beta-cell mass undergoes during the different stages of disease progression. Reformed islets allow a non-invasive, high-resolution system to quantitatively image at the single-islet level which can be useful for the evaluation of islet viability, proliferation, and the effects of novel anti-diabetic compounds. For example, reformed islets can be used to report macrophage infiltration by coculturing with transgenic-labelled fluorescent macrophages.

In summary, we have refined a method for generating a static reformed islet platform which can be applied to both mouse and human cells and cultured over the long-term. Reformed islets are genotypically, phenotypically, and functionally like the primary islets they are derived from and importantly harbour a resident immune population, which is missing in other platforms. These properties make reformed islets highly amenable to exploring islet cell-cell communication for example using multiple imaging modalities and repeated functional assessment. The platform could easily be optimised for use in a multi-well format for high content screening programmes and will likely prove to be a relatively easy platform to genetically manipulate given the dispersion step in the methodology. Dispersed islets cells are far easier to transfect whether using lipid-based transfection reagents, viral vectors, or electroporation [55, 56]. We hope the platform will be a useful tool to further our understanding of diabetes pathology.

---

Supplemental Table 1: Primary Antibodies.

**Supplemental Table 1: Primary Antibodies**
| Target | Company | Product # | Concentration |
| --- | --- | --- | --- |
| Insulin | Abcam | ab7842 | 1:200 |
| Glucagon | Merck Life Science UK Limited | G2654 | 1:1000 |
| Somatostatin | Abcam | ab30788-125ul | 1:50 |
| Ki67 | Abcam | ab15580 | 1:200 |
| CD80 | Bio-Techne | AF740-SP | 1:100 |
| IBA1 | Alpha Laboratories Limited | 019-19741 | 1:200 |
| CD45 | Abcam | ab40763 | 1:100 |
| CD8α | Cell Signaling | 98941S | 1:100 |
| CD4 | BD Pharmingen™ | 557667 | 1:50 |

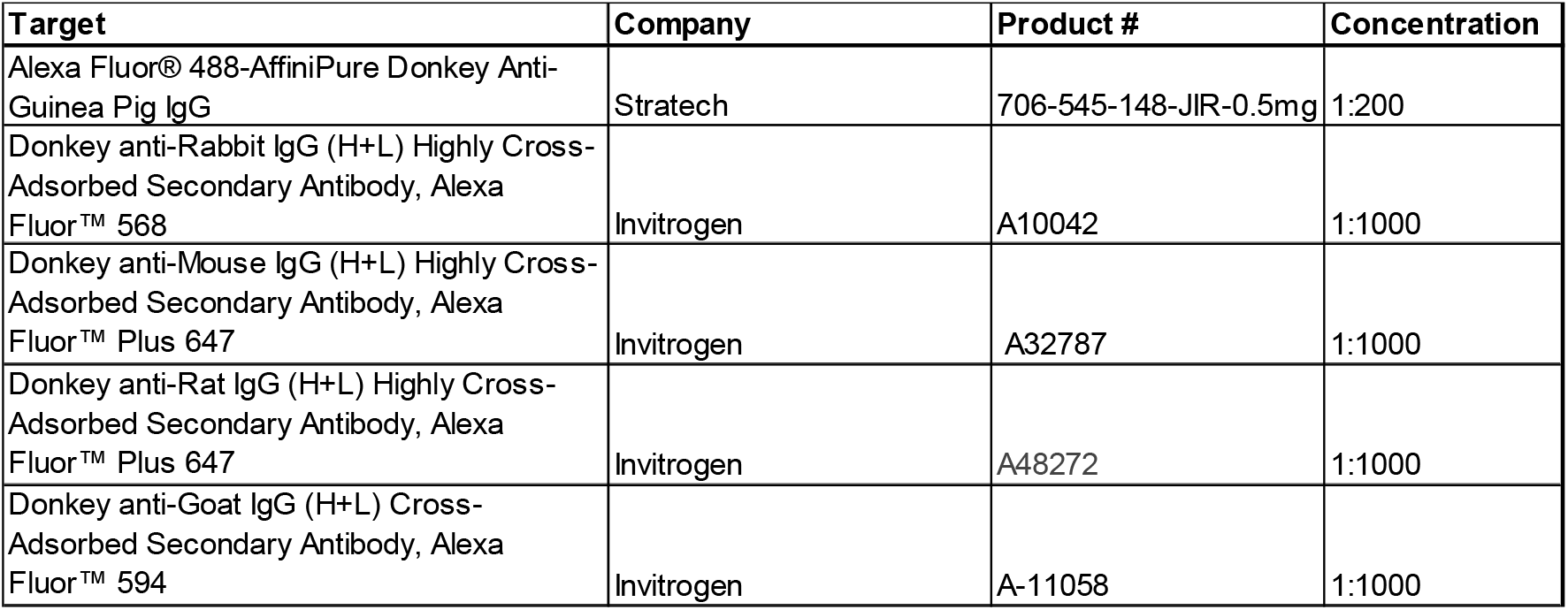
Supplemental Table 2: Secondary Antibodies.

## Figure legends

**Figure S1.**
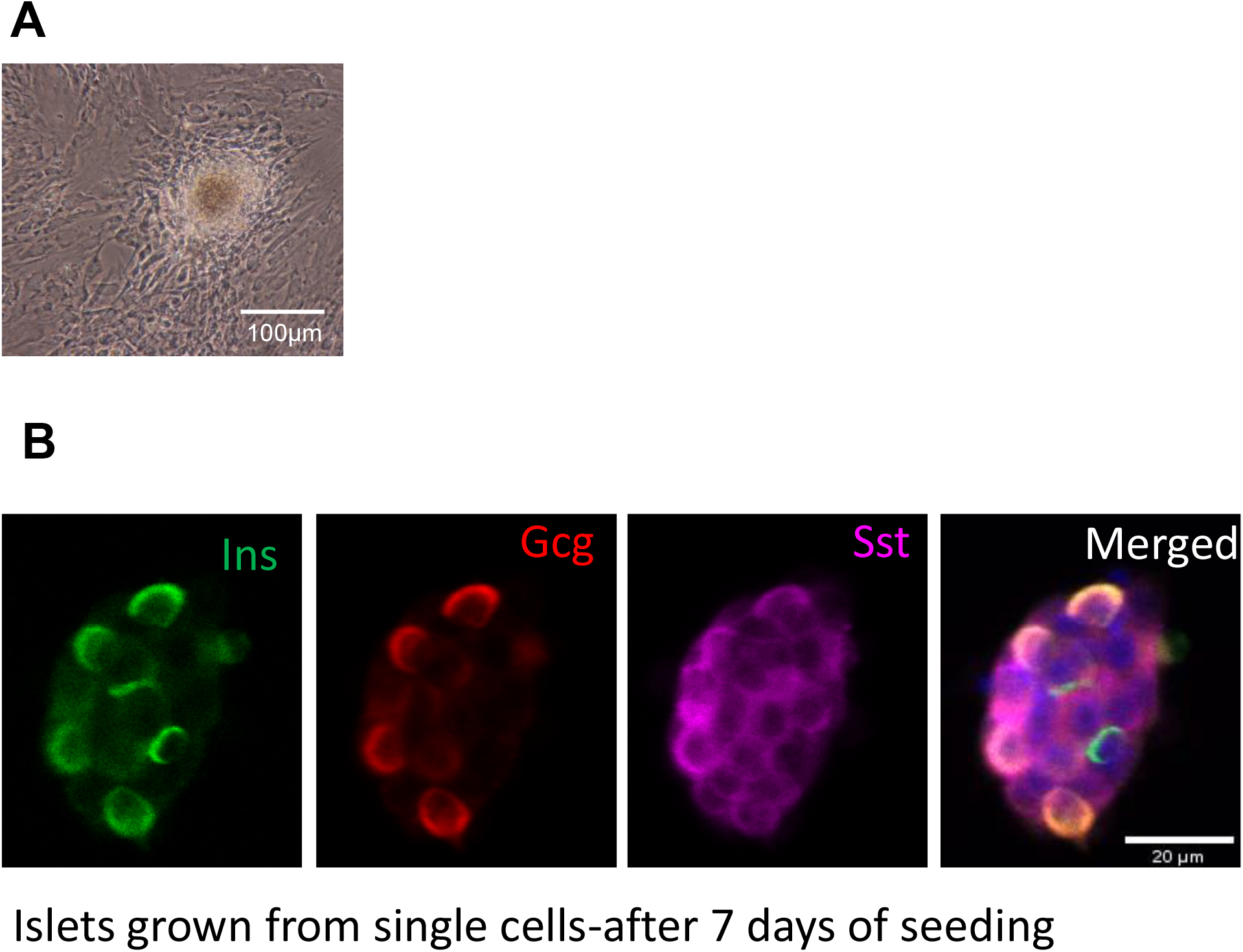
Light microscopy images show that human islet cells formed 3-dimensional spheroid structures after 7 days of seeding. Scale bar is 100µm (A). Immunofluorescence detection by confocal microscopy of islet hormones in maturing mouse reformed at day 7 confirmed that developing reformed islets expressed insulin (Ins; green), glucagon (Gcg; red) and somatostatin (Sst; purple). Scale bar is 20µm.

**Figure S2.**
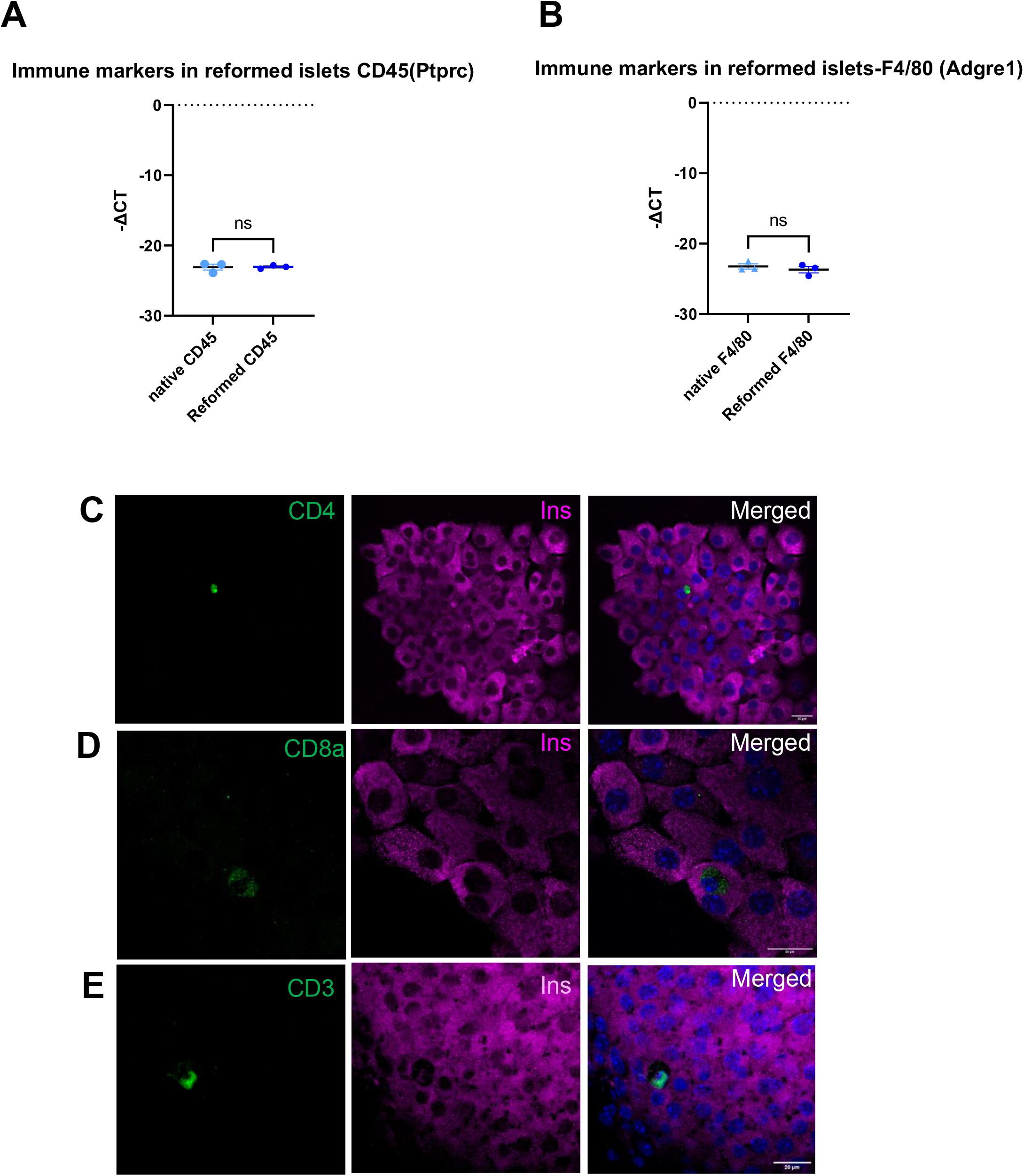
Immune population in reformed islets. (A, B) Delta Ct values distribution graphs of Ptprc (CD45, all immune cell marker) and Adgre1(Macrophage marker) in native and reformed mouse islets detected by quantitative PCR. Immunostaining of the reformed islets with antibodies directed against insulin (Ins; purple) and immune markers CD4 (C), CD8a (D) and CD3 (F) (green) confirmed the presence of both resident macrophages and resident T-cells in reformed islets. Scale bar is 20µm.

**Figure S3.**
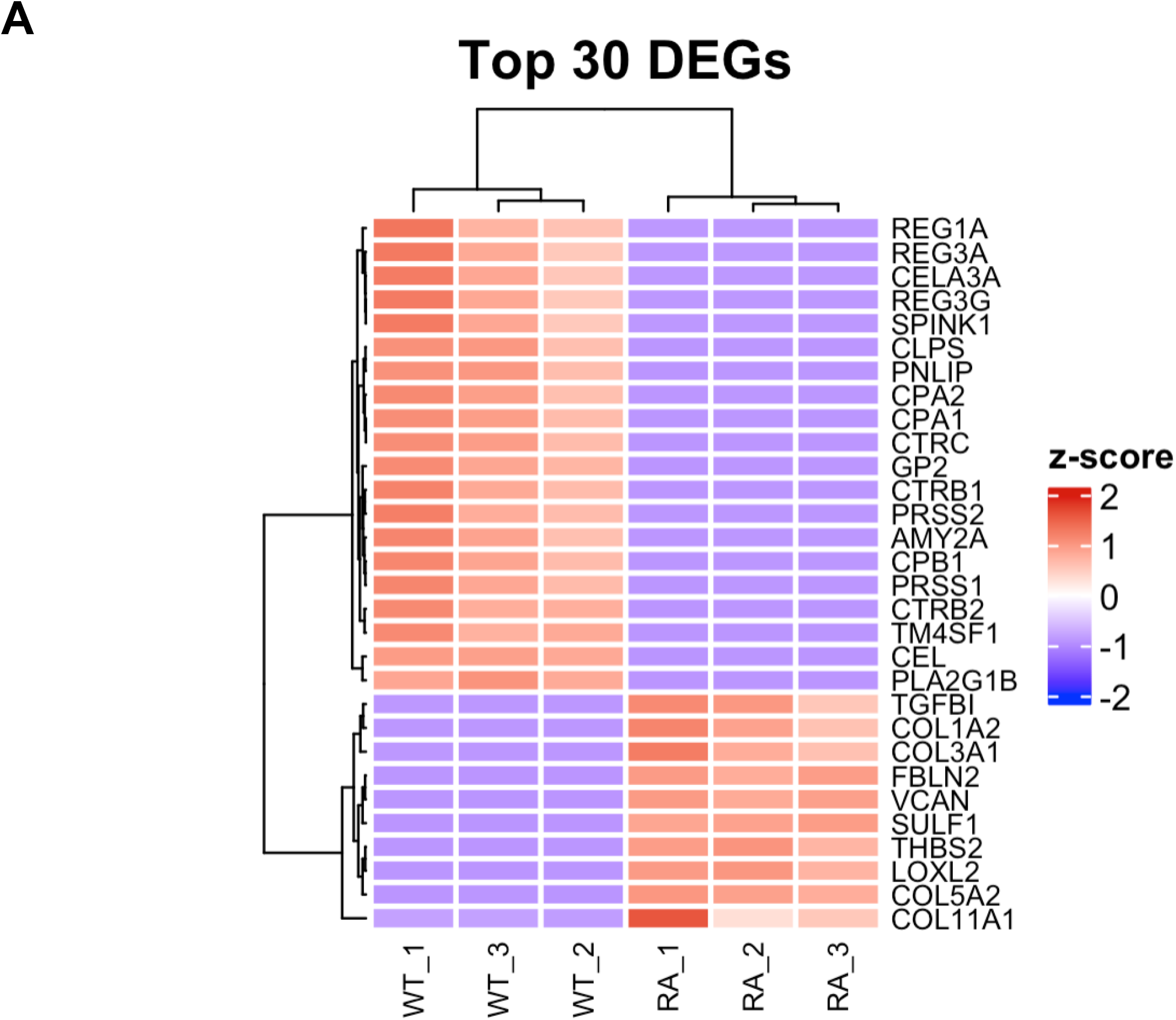
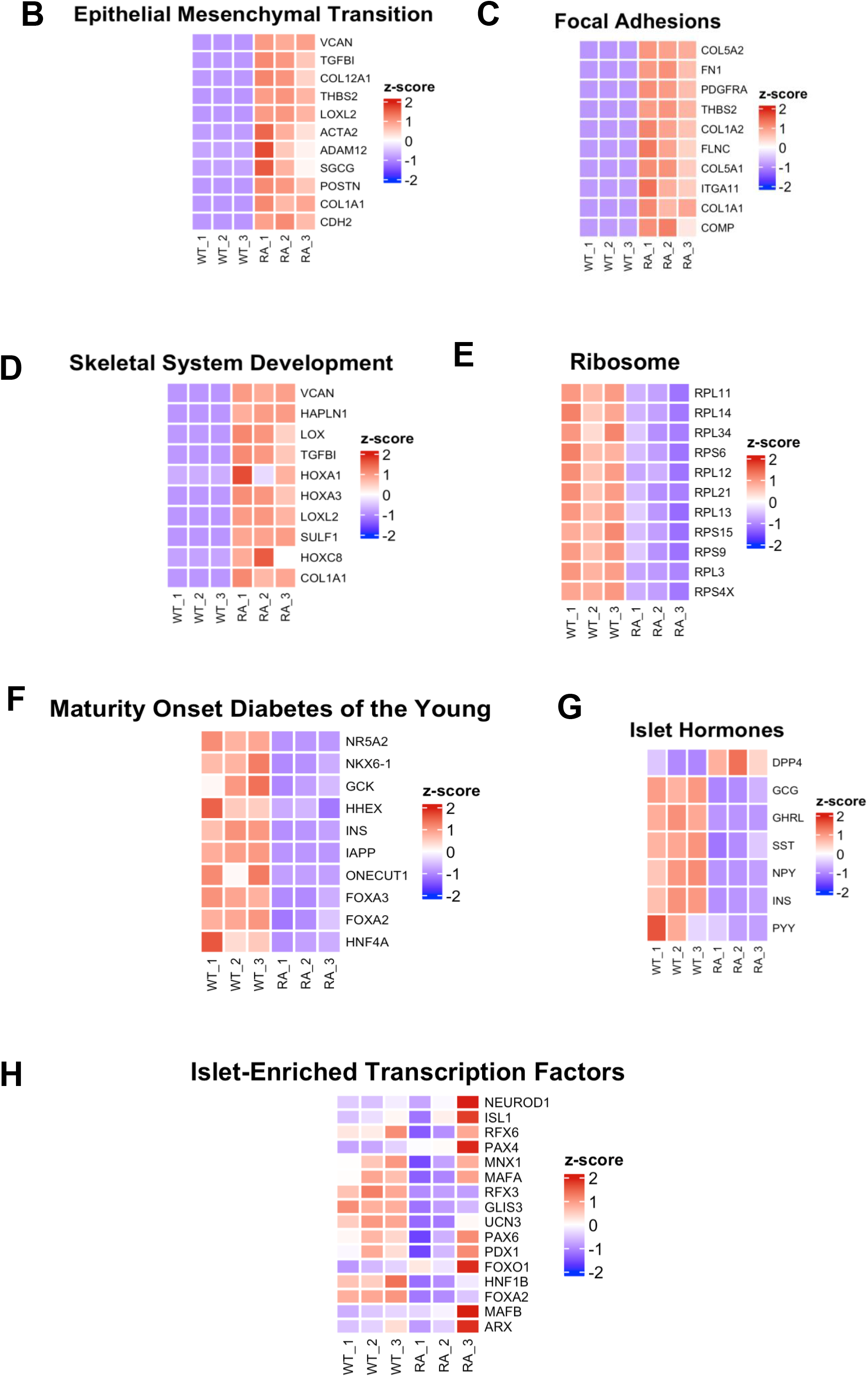
Transcriptomic analyses of native and reformed human islets. (A) Heatmap illustrating top 30 differentially expressed genes in reformed islets. (B, C, D) Heatmaps illustrating genes differentially enriched in EMT, Focal adhesion and Skeletal development pathway process. (E) KEGG enriched pathways related to ribosome biogenesis. (F) Pathways enriched in Maturity Onset of the Diabetes of the Young. (G) Heatmap of mRNA expression of islets hormones. (H) Heatmap of selected subset of islet-enriched transcription factors between native and reformed islets.

## AUTHOR CONTRIBUTIONS

NH and GAB designed and managed the studies. All authors contributed to the writing and editing of the manuscript. NH, KT, MZ, YL carried out all the islet studies. JP isolated and provided the T-cells. MW, MJ, GAB and TP analysed the sequencing data.

## Notes

### Competing Interest Statement

The authors have declared no competing interest.

### Summary of Updates

Added Naila Haq as corresponding author Changed spelling of Min Zhao

